# Microbiota-derived secondary bile acids promote STING activation and antitumor activity

**DOI:** 10.1101/2025.04.16.649255

**Authors:** Xinglin Yang, Xing Zhang, Weichao Li, Anant Gharpure, Althea Hansel-Harris, Yan-Zhu Hsieh, Stanislav Dikiy, Andreas F. Tillack, Chongyang Wu, Ariana Sulpizio, Kyong T. Fam, Xiaohui Zhao, Stefano Forli, Luke L. Lairson, Andrew B. Ward, Christopher G. Parker, Howard C. Hang

## Abstract

Microbiota metabolism generates diverse bile acids that are associated with health and disease, but the molecular targets and mechanisms of action for these metabolites have not been fully elucidated. Using deoxycholic acid (DCA) photoaffinity probes, with cholic acid (CA) probe as a comparator, we found many protein targets of microbiota-derived DCA in mammalian cells. Of note, we discovered that DCA binds the transmembrane domain of stimulator of interferon genes (STING), and enhances agonist-induced accumulation of higher-order STING species, type I interferon signaling and antitumor immunity. Moreover, DCA-producing microbiota species enhanced STING agonist-induced antitumor immunity primarily through the activation of natural killer (NK) cells. This reverse chemical microbiology approach revealed an unpredicted mechanism of action for DCA on STING activation and suggests that specific secondary bile acids and their associated microbiota species may promote NK activation and impact the efficacy of STING-targeted therapeutics.

## Main Text

Bile acids are sterol metabolites that are associated with diverse phenotypes in health, disease and therapy responses(*1*), but their specific mechanisms of action are still enigmatic. Biosynthesis of bile acids starts with oxidation and hydroxylation of cholesterol in the liver followed by conjugation with glycine or taurine to generate primary bile acids(*1*). Export of primary bile acids into the intestine facilitates cholesterol turnover and disposal as well as aids in digestion. In the gut, primary bile acids are deconjugated, oxidized and epimerized by specific microbiota species to generate a variety of secondary bile acids(*2–5*) (**Fig. 1a**). In recent years, these structurally diverse microbiota-derived bile acids have emerged as signaling molecules that regulate a variety of physiological processes, ranging from direct antibacterial activity(*6, 7*) to modulation of host metabolism(*8–11*), immunity(*12, 13*), viral infection(*14, 15*), vaccine responses(*16*), aging(*7, 17*) and tumor progression (*18–22*). Of note, bile acids were discovered as endogenous ligands for farnesoid X receptor (FXR), which regulates the expression of target genes involved in hepatic bile acid synthesis, secretion and intestinal reabsorption(*23, 24*) and are also implicated in T cell-driven inflammation(*13*). In addition, the G protein-coupled receptor TGR5 has been reported to sense bile acids and regulate intestinal homeostasis and inflammation(*12, 25*). Moreover, secondary bile acids have been reported to bind and regulate the activity of specific nuclear hormone receptors like RORγt and NR4A1 in T cells to control their differentiation and activation(*26–30*). Secondary bile acids have also been shown to phenocopy anti-aging effects of calorie restriction by binding TUB-like protein 3(*17, 31*). Metabolomic studies indicate that hundreds of additional conjugated and unconjugated bile acids can be found in different mammalian tissues(*4, 9*), which necessitates further mechanistic and functional studies.

**Figure 1.**
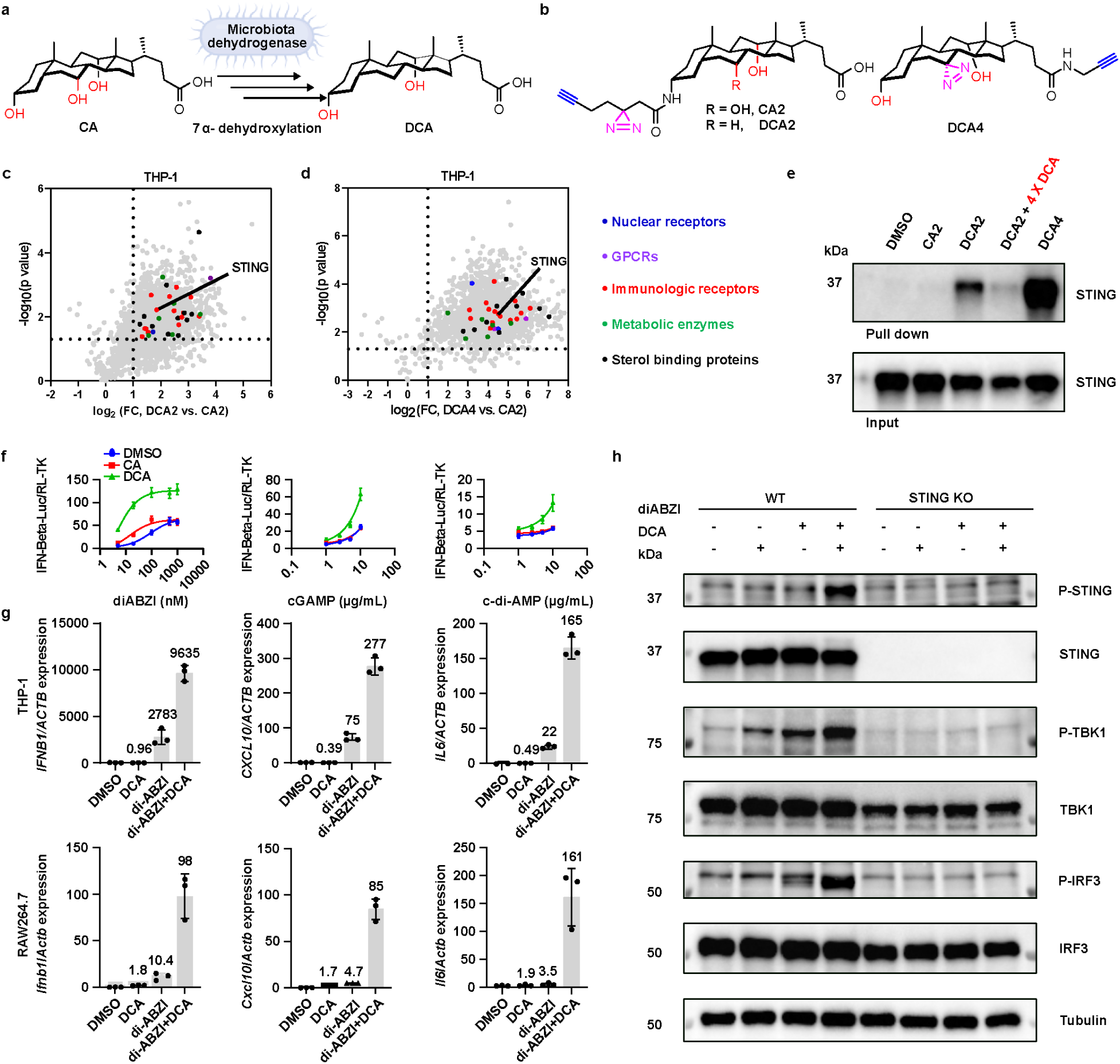
Microbiota-derived DCA interacts with STING and potentiates STING agonist-induced innate immune signaling. (a) Gut microbiota convert cholic acid (CA) into deoxycholic acid (DCA) through a multistep 7α-dehydroxylation pathway. This pathway is largely encoded by the bile acid-inducible *bai* operon, with related bile acid oxidoreductases contributing to intermediate oxidation and reduction steps. (b) Chemical structure of bile acid photoaffinity probes. (c, d) DCA2 (25 µM) and DCA4 (25 µM) bile acid photoaffinity reporters enrich more proteins compared to CA2 in THP-1 cells. STING is one of the proteins that are selectively enriched by DCA photoaffinity reporters. Other DCA-enriched proteins are highlighted by different colors. (e) Validation of bile acid-STING interaction in THP-1 by reverse western blot. THP-1 cells were treated with bile acid photoaffinity reporters (25 µM) with or without excess of DCA for 0.5 h, followed by 365-nm light irradiation. The labeled proteins were reacted with az-biotin and enriched with streptavidin beads (‘pulldown’). Equal loading was validated by immunoblotting of input samples before enrichment (‘input’). (f) HEK293T transfected with pMSCV-hygro-STING, IFN-Beta_pGL3 (a luciferase reporter plasmid under control of the IFN-β promoter) and pRL-TK control plasmid were treated with different doses of STING agonists (diABZI, cGAMP, c-di-AMP) in the presence or absence of DCA and CA (50 µM) (For diABZI, *n* = 5, for cGAMP and c-diAMP, *n* = 3, biologically independent samples). (g) RT-qPCR analysis of *IFNB1, CXCL10, and IL6* in human THP-1 monocytes and their murine orthologs *Ifnb1, Cxcl10, and Il6* in RAW264.7 macrophages. Cells were treated with diABZI (50 nM for THP-1, 200 nM for RAW264.7) and DCA (50 µM) alone or together for 4 hours (*n* = 3 biologically independent samples). (h) Western blot analysis of cell lysates from THP-1 cells (WT and STING KO) treated with diABZI (50 nM) and DCA (50 µM) alone or together for 4 hours.

To address the direct protein targets and mechanisms of action for different bile acids, we and others have generated photoaffinity probes to identify and validate their protein targets in bacteria(*32–35*) and mammalian cells(*31, 36*). These bile acid photoaffinity probes contain a photoreactive diazirine group and an alkyne handle, allowing UV-induced covalent crosslinking to nearby interacting proteins in cells and subsequent detection using bioorthogonal chemistry(*37, 38*). In this study, we explored the protein targets of the secondary bile acid deoxycholic acid (DCA) compared to its primary bile acid precursor, cholic acid (CA), in monocytes using photoaffinity probes and discovered that DCA binds to the transmembrane domain of stimulator of interferon genes (STING) and allosterically promotes its activation in combination with native and synthetic agonists. Notably, DCA enhanced STING agonist induction of type I interferon signaling and antitumor immunity. Moreover, DCA-producing microbiota species such as *Clostridium scindens* also enhanced STING agonist antitumor immunity. Our immune profiling of tumor-infiltrated lymphocytes suggests DCA enhances STING agonist antitumor immunity primarily through the activation of NK cells. These studies suggest specific bile acids and their associated microbiota species can modulate host immunity and efficacy of immunotherapies.

### Chemoproteomics reveals DCA binding to STING

To explore the bile acid protein targets in mammalian cells, we focused on DCA, a secondary bile acid generated by the gut microbiota dehydroxylation of CA (**Fig. 1a**)(*39*). The removal of a hydroxyl group at the C7 of the sterol significantly alters the physicochemical properties of the bile acid, potentially conferring unique membrane penetrating and protein binding capabilities of DCA. DCA has been ascribed both protective and pathogenic activities on host physiology. On the protective side, microbiota-derived secondary bile acids, including DCA, have been implicated in colonization resistance against enteric pathogens, modulation of host metabolic and immune responses and restriction of viral dissemination through bile acid–type I interferon signaling axes(*6, 14–16*). These findings suggest that DCA can contribute to beneficial host–microbiota interactions under specific physiological or infectious contexts. Conversely, elevated or chronic exposure to DCA and related secondary bile acids has been associated with pathogenic outcomes, including tissue injury, inflammation as well as enhanced tumor progression in the liver and colorectal cancer models(*18–21*). DCA has also been developed as a lipolytic therapeutic agent, reflecting its potent membrane-active properties at high local concentrations(*40*). Despite these notable associations with host physiology, immunity and therapy, the mechanisms of action for DCA on specific proteins have not been well-characterized.

To explore the immunomodulatory activities of DCA, we performed chemoproteomic studies in human THP-1 monocytes. Since chemical modifications can influence metabolite protein binding, we designed two DCA photoaffinity probes with distinct modification sites to improve target coverage and reduce probe-specific bias. DCA2 introduces the photocrosslinking group from the 3-hydroxyl position while largely preserving the steroid nucleus and carboxylate-containing side chain. Alternatively, DCA4 incorporates a diazirine into the steroid nucleus and introduces the alkyne handle through an amide linkage at the carboxylate side chain. These complementary probes allowed us to capture a broader set of DCA-interacting proteins and prioritize candidate targets for subsequent investigation. Using different photoaffinity probes for DCA and CA (**Fig. 1b, Extended Data Fig. 1a**), our quantitative comparative chemoproteomic analysis of THP-1 monocytes revealed many candidate DCA-selective protein targets (**Fig. 1c, d**), including sterol metabolic enzymes, membrane proteins, nuclear receptors and immunity-associated proteins (**Extended Data Fig. 1b, Tables S1-3, supplementary Data S1**). Competition with parent bile acid was used to help distinguish specific bile acid-interacting proteins from nonspecific probe-enriched proteins (**Extended Data Fig. 1b**). Notably, STING (TMEM173) was one of the DCA probe-selective protein targets compared to CA probe (**Fig. 1c, d, Extended Data Fig. 1b**). Western blot analysis of photoaffinity probe-enriched proteins showed that STING was crosslinked by the DCA probes, but not the CA probe, which was significantly reduced when THP-1 cells were co-incubated with DCA (**Fig. 1e**). DCA also reduced photoaffinity probe crosslinking of FXR and STING endogenously expressed in HT-29 cells (**Extended Data Fig. 1c**) and human STING overexpressed in HEK293T cells (**Extended Data Fig. 1d**). These results suggest that DCA may directly bind STING in mammalian cells.

STING is an important innate immune receptor that is activated by cyclic dinucleotides (CDNs), including cyclic guanosine monophosphate–adenosine monophosphate (cGAMP) synthesized by cyclic GMP-AMP synthase (cGAS) in response to cytosolic DNA(*41–43*). Upon CDN binding, endoplasmic reticulum (ER)-localized STING oligomerizes and traffics through ER–Golgi and endolysosomal compartments, where it recruits TANK-binding kinase 1 (TBK1). TBK1 phosphorylates itself, STING and transcription factors such as interferon regulatory factor 3 (IRF3), leading to induction of type I interferons (IFNs), NF-κB-dependent cytokines and other immune mediators that promote host defense against infections and tumors(*44–46*). Based on these properties, synthetic STING agonists have been developed as therapeutic leads for infectious diseases and cancer(*29, 47–49*). Compounds that mimic CDNs, such as diABZI, SR-717 and MSA-2, target the cyclic dinucleotide-binding domain of STING and potently induce type I IFN responses(*50–52*). In addition, small molecules such as C53, NVS-STG2 and Y-320 engage the STING transmembrane region and modulate STING oligomerization and immune activation(*53–55*). Representative compounds targeting the CDN-binding domain and transmembrane domain of STING are shown in **Extended Data Fig. 2a, b**.

Recent studies further indicate that STING activation is highly sensitive to its membrane lipid environment. Cholesterol-binding motifs in STING have been shown to regulate ER retention and STING oligomerization(*56, 57*). More recently, PtdIns(3,5)P₂ was identified as an endogenous STING ligand that promotes agonist-dependent STING activation and acts together with cholesterol to stabilize higher-order STING oligomers and enable TBK1 activation(*58, 59*). In parallel, the Golgi-resident lipid PI4P and the chemical agonist GNE-6468 were reported to cooperatively activate STING by inducing transmembrane helix rearrangement(*60*). These findings suggest that the STING transmembrane and membrane-proximal regions function as lipid- and small molecule-sensitive domains. Moreover, microbiota-derived metabolites, including bacterial CDNs, can modulate STING signaling and shape type I interferon-dependent immune responses(*61–64*). However, whether microbiota-derived bile acids such as DCA can directly bind STING and impact STING activation was unknown.

### Bile acids promote STING activation *ex vivo*

We therefore investigated if DCA alone or in combination with STING agonists modulated type I IFN responses. Using an IFN-β luciferase reporter assay, we found that DCA but not CA enhanced the activity of CDNs (cGAMP and c-di-AMP) and synthetic STING agonists such as diABZI (**Fig. 1f, Extended Data Fig. 2e**). DCA alone did not significantly activate STING at concentrations up to 200 μM, however, it dose-dependently enhanced diABZI-induced STING phosphorylation in THP-1 cells (**Extended Data Fig. 2c, d**). To further investigate the STING activation, we measured *IFNB1, CXCL10, and IL6* in human THP-1 monocytes and their murine orthologs *Ifnb1, Cxcl10, and Il6* in RAW264.7 macrophages using RT-qPCR (**Fig. 1g**). Co-treatment with DCA and diABZI induced significantly higher levels of these cytokines compared to either treatment alone (**Fig. 1g**). Treatment of THP-1 monocytes with DCA and diABZI increased the phosphorylation of STING, TBK1, and IRF3 compared to either compound alone, which was also abrogated in STING knockout (KO) cells (**Fig. 1h**).

To further characterize the scope and specificity of DCA-STING activation, we transfected differentiated THP-1 monocytes with high molecular weight DNA (HT-DNA), which activates cGAS to produce endogenous cGAMP and observed DCA enhancement of HT-DNA-induced STING activation (**Extended Data Fig. 3a**). DCA also enhanced the activity of other synthetic STING agonists (SR-717, MSA-2 and DMXAA) in HEK293T cells expressing human STING and an IFN-β reporter (**Extended Data Fig. 3b**) as well as other STING variants, including murine STING, STING-HAQ (R71H, G230A and R293Q), and STING-H232 (R232H)(*65*) (**Extended Data Fig. 3c**). DCA enhancement of STING activation was also observed with CDNs (**Extended Data Fig. 2g, h**) and diABZI in primary human peripheral blood mononuclear cells (PBMCs) (**Extended Data Fig. 2f, i, j**) as well as differentiated THP-1 (**Extended Data Fig. 3d**) and HT-29 cells (**Extended Data Fig. 3e**). We also observed more STING puncta formation (**Extended Data Fig. 4a,b**) and oligomerization (**Extended Data Fig. 4c**) in HeLa cells that were co-treated with diABZI and DCA, but not CA. Interestingly, pre-treatment of THP-1 monocytes with diABZI enhanced the interaction between the DCA probe and STING (**Extended Data Fig. 4d,e**). This enhanced interaction was suppressed by BB-Cl (**Extended Data Fig. 4d,e**), an inhibitor of STING activation (**Extended Data Fig. 4f**)(*66*).

### Bile acids bind STING dimer-dimer interface

To investigate the binding site of DCA on STING, we conducted competitive ligand photoaffinity crosslinking studies in cells and structural studies *in vitro*. Notably, NVS-STG2, a synthetic molecule reported to target the STING dimer-dimer interface reduced DCA probe photocrosslinking with STING in cells (**Fig. 2a,b**). In contrast, diABZI which binds the CDN-binding domain failed to affect the DCA probe-STING photocrosslinking, even with 10-fold excess (**Extended Data Fig. 4g**). To elucidate the DCA-STING binding site, we employed single-particle cryo-electron microscopy (cryo-EM). We included STING agonists cGAMP and C53 to capture the ligand-activated protein state and stabilize potential oligomers, similar to previous studies with NVS-STG2. From these studies, we obtained a 2.9 Å resolution map of a STING tetramer complexed with ligands (cGAMP, C53 and DCA) (**Extended Data Fig. 5 and Table S4**). Notably, we observed additional molecular density between the transmembrane domains of STING dimers (**Fig. 2c**), which was absent in the STING-cGAMP-C53 structure(*53*), suggesting this region as a potential binding site for secondary bile acids like DCA.

**Figure 2.**
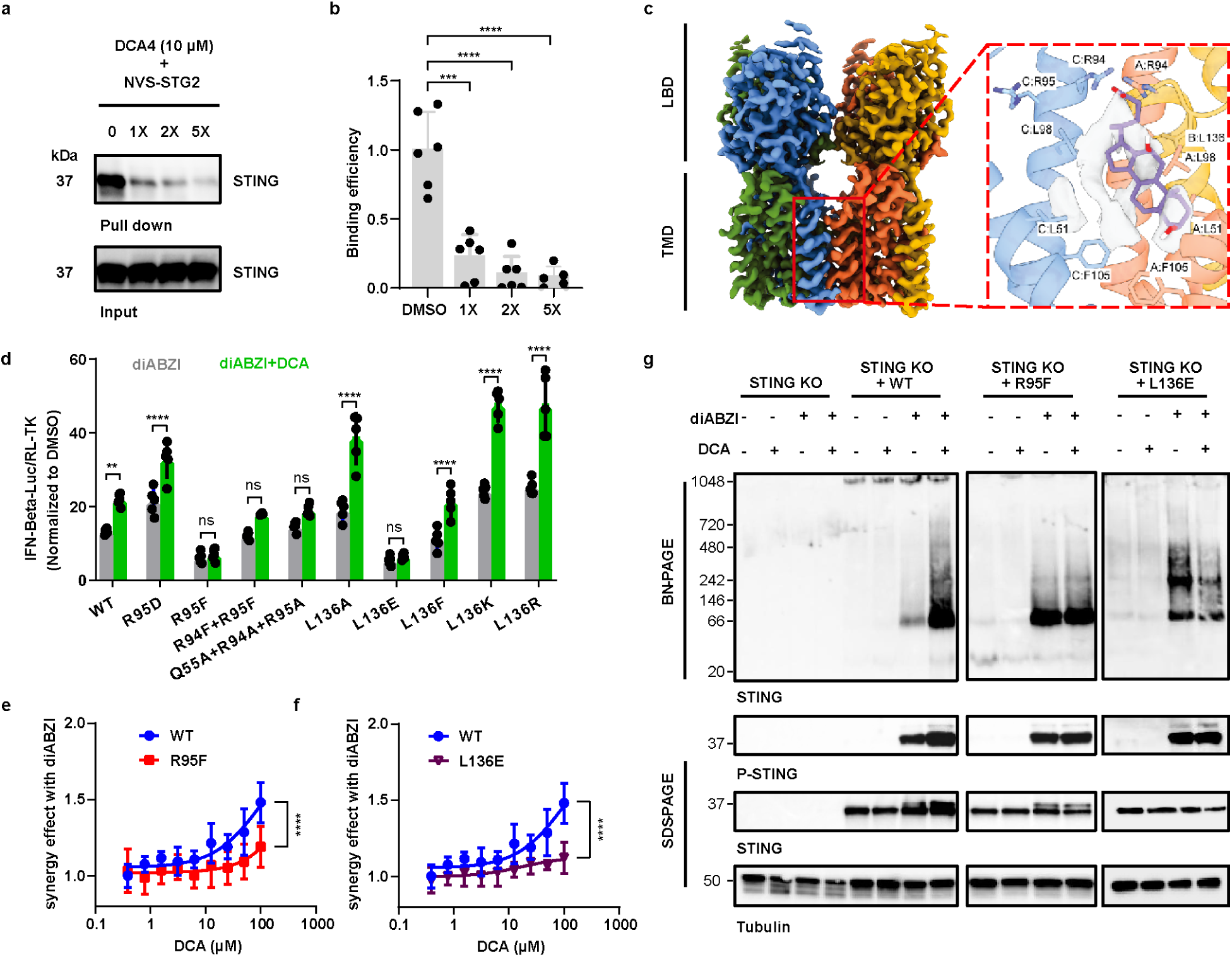
Bile acids target at STING dimer-dimer interface. (a) HEK293T transfected with pMSCV-hygro-STING were treated with DCA4 (10 µM) in the presence of different concentrations of NVS-STG2 for 30 min at 37 °C, followed by 365-nm light irradiation. The labeled proteins in cell lysis were reacted with az-biotin and enriched with streptavidin beads (‘pulldown’). Equal loading was validated by immunoblotting of input samples before enrichment (‘input’). (b) Quantification of binding efficiency in A by grayscale analysis (pull down/input) (*n* = 5 - 6 biologically independent samples.). Data were analyzed by unpaired two-sided t-test. (c) High-resolution cryo-EM map of the two STING dimers with highlight of dimer-dimer interface molecular density. Docked model of DCA in STING tetramer. LBD: ligand binding domain, TMD: transmembrane domain. (d-f) HEK293T transfected with pMSCV-hygro-STING (WT and mutations), IFN-Beta_pGL3 and pRL-TK control plasmid were treated with diABZI (20 nM) in the presence or absence of DCA (*n* = 5 biologically independent samples). A two-way ANOVA followed by Sidak’s multiple comparisons test was used to calculate adjusted P values. *P < 0.05, **P < 0.01, ***P < 0.001, ****P < 0.0001. (g) STING oligomerization induced by DCA, diABZI or both. THP1-Dual STING KO cells (STING-WT, STING R95F, STING L136E stable expression cell lines) were stimulated with DCA in the presence or absence of diABZI for 2 hours. STING oligomerization was analyzed by Blue native PAGE (BN-PAGE), and the indicated proteins were detected by immunoblotting.

To explore whether DCA could occupy the STING dimer–dimer interface, we used a custom Monte Carlo docking engine. Affinity grid maps were calculated with the AutoDock4.2 force field and modified using our CryoXKit(*67*) software to include guidance from the C2-symmetric cryo-EM density. The resulting docked model is shown in **Extended Data Fig. 6a**, where we have modeled one bile acid molecule into one half of the C2 map. The carboxylate of DCA is predicted to form strong charge-charge interactions with R94 of both the A (red) and C (blue) STING monomers, vertically orienting the ligand parallel to the surrounding helices. The rest of the molecule forms interactions mostly with the A and B (yellow) monomers. This single, static binding pose only explains a limited portion of the observed ligand density. Therefore, we generated a Boltzmann-weighted map of all poses sampled during the docking simulation, to better represent the full volume sampled during the run and allow for more direct comparison to the cryo-EM density. Notably, this map also includes the sampled positions of oxygens and polar hydrogens that may be poorly resolved in the corresponding cryo-EM density. The docking Boltzmann map (outer purple surface in **Extended Data Fig. 6a**) shows that the docking does largely sample all the cryo-EM density volumes assigned to the ligand, including those not explained by the single predicted binding pose. These results suggest that DCA may bind at the STING transmembrane dimer-dimer interface to enhance oligomerization and promote agonist-induced activation.

To further assess the evolutionary conservation of the proposed DCA-associated region, we aligned representative vertebrate STING orthologs. This analysis showed that the putative DCA-associated region lies within a broadly conserved transmembrane region of STING. Several hydrophobic residues that contribute to the transmembrane dimer–dimer interface are conserved or maintain hydrophobic character across species, whereas residues near the predicted DCA carboxylate, including R94 and R95 in human STING, show species-dependent variation (**Extended Data Fig. 5h**). These results suggest that DCA engages a partially conserved transmembrane regulatory interface with species-variable electrostatic features, rather than a strictly conserved DCA-specific pocket.

To validate the proposed bile acid binding site, we generated STING transmembrane mutants and assessed their response to diABZI and DCA. Several mutants, such as L136A, L136K, and L136R, showed increased sensitivity to DCA activation in HEK293T cells (**Fig. 2d**). In contrast, other STING mutants, including R95F, the R94F/R95F double mutant as well as L136E, exhibited reduced sensitivity to DCA-induced enhancement of STING activation in HEK293T cells (**Fig. 2d**). R95F and L136E were selected for further analysis because these single substitutions attenuated DCA-mediated enhancement while retaining sufficient responsiveness to diABZI to permit evaluation of the DCA-dependent effect. Expression of these STING mutants R95F and L136E in HEK293T (**Fig. 2e,f**) or STING-KO THP-1 cells (**Fig. 2g**), which are reported to express other bile acid receptors such as FXR(*68*) and TGR5(*15*), still showed attenuated DCA enhancement of diABZI-induced STING activation. Additional oligomerization analysis of the Q55A/R94A/R95A mutant is shown in **Extended Data Fig. 6b**. These mutagenesis and activity data support the model that DCA binds directly to the STING transmembrane domain dimer-dimer interface to modulate immune signaling.

To assess the specificity of BAs on STING activation, we compared the effects of different conjugated and free bile acids. DCA consistently exhibited the most potent activity among all tested BAs (**Extended Data Fig. 6c**). Structural modifications at the 3-hydroxyl position, such as isomerization or oxidation, significantly reduced DCA enhancement of STING activation (**Extended Data Fig. 6c**). Interestingly, alloLCA, but not LCA or isoLCA, showed comparable levels of enhanced STING activation as DCA (**Extended Data Fig. 6c**). These results demonstrate DCA, amongst the BAs tested, most effectively enhances STING activation in a variety of different cell types. Molecular docking of other bile acids (**Extended Data Fig. 6d**) shows the inactive sterols (CA, iso-DCA, UDCA and iso-LCA) do not make ideal contacts with STING transmembrane dimer-dimer interface compared to other active secondary bile acids such as alloLCA (**Extended Data Fig. 6d**), which adopts similar conformation to DCA (**Extended Data Fig. 6a**). These comparative bile acid analyses suggest that DCA-mediated STING modulation depends on multiple structural features. The reduced activity of CA relative to DCA indicates that the C7 hydroxylation pattern influences activity, whereas loss of activity upon 3-hydroxyl isomerization or oxidation suggests that the orientation and hydrogen-bonding properties of this group are important. The activity of alloLCA, but not LCA or isoLCA, further suggests that steroid ring configuration affects whether bile acids can adopt a productive binding pose in the STING transmembrane pocket. Finally, the predicted interaction between the DCA side-chain carboxylate and R94 suggests that side-chain positioning may help orient bile acids at the dimer–dimer interface. Thus, hydroxylation pattern, steroid configuration, and side-chain positioning may collectively influence DCA binding to STING and enhancement of STING activation. Together, these findings indicate that secondary bile acids such as DCA bind to a large pocket at the STING transmembrane dimer-dimer interface to enhance its ligand-induced oligomerization and activation of downstream signaling.

### DCA boosts STING agonist antitumor immunity

As STING activation has been shown to promote antitumor immunity(*48, 49*), we evaluated the bile acids with diABZI in tumor models. For these studies, we employed intratumoral (IT) injection of the compounds (**Fig. 3a**) in subcutaneous tumor models (*69*) and utilized a lower dose of diABZI based on pilot dose-ranging experiments (**Extended Data Fig. 7a**). This diABZI dose alone produced minimal antitumor activity, thereby enabling assessment of potential synergistic effects with bile acids. IT injection of DCA (2.5 mg/kg) combined with a low dose of diABZI (50 μg/kg) significantly reduced tumor size, whereas neither treatment alone showed an effect (**Fig. 3b, Extended Data Fig. 7b**). Notably, this effect was DCA-specific, as CA together with diABZI showed no antitumor activity against B16-F10 melanoma tumors (**Fig. 3c**). We next investigated whether DCA synergized with STING agonist treatment in additional tumor models beyond B16-F10 melanoma. Subcutaneous tumors were established using less immunogenic Panc02 pancreatic ductal adenocarcinoma cells (*70*) and immunogenic MC38 colorectal carcinoma cells. Similar to B16-F10, low-dose diABZI alone failed to significantly reduce Panc02 tumor growth and required combination with DCA to achieve antitumor effects (**Extended Data Fig. 7c**). In contrast, diABZI alone was sufficient to reduce MC38 tumor burden (**Extended Data Fig. 7d**), consistent with previous studies that MC38 tumors are more responsive to immune checkpoint blockade and STING agonist treatment(*71*).

**Figure 3.**
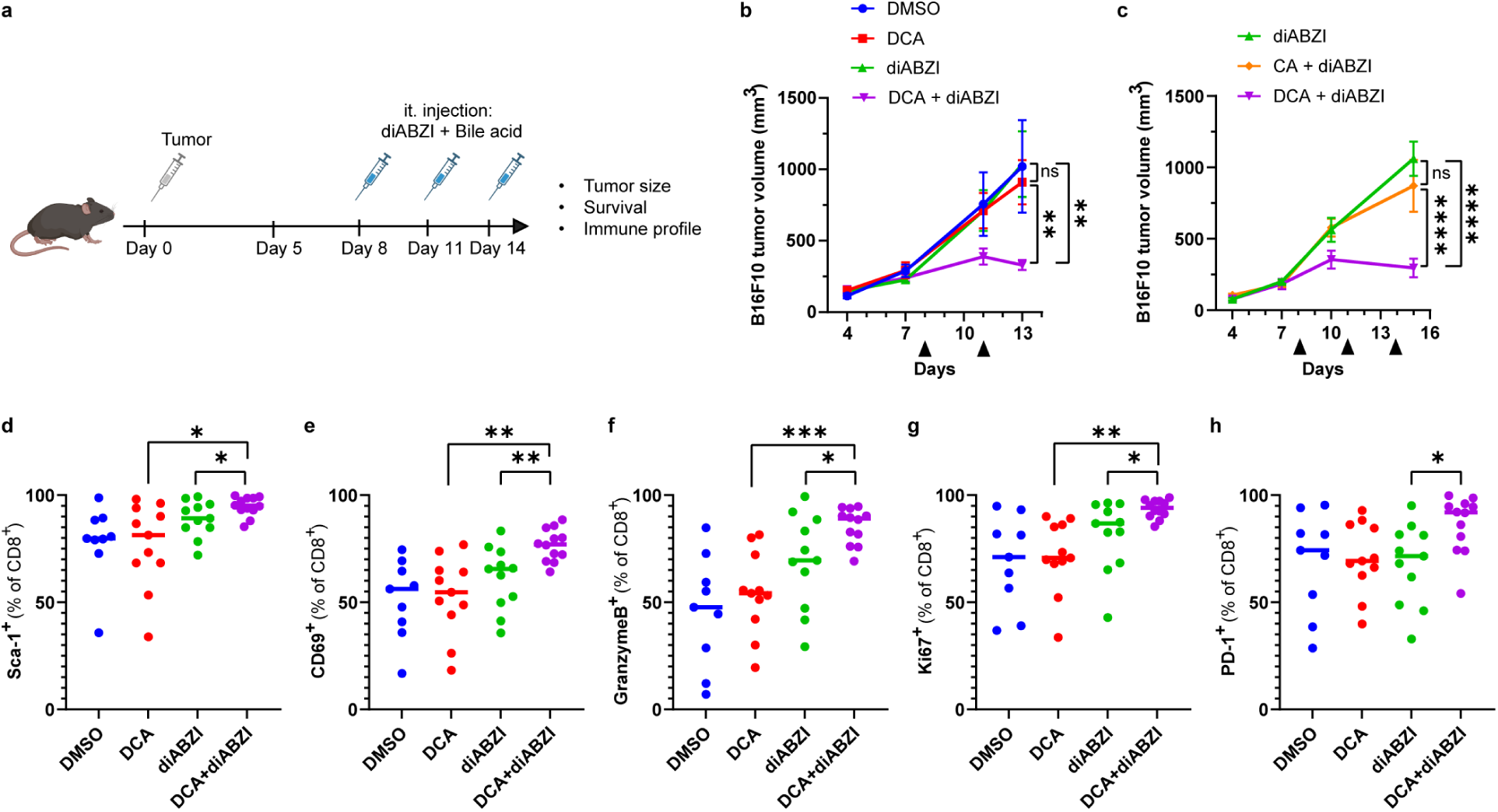
DCA boosts STING-targeted antitumor immunity *in vivo*. (a) Schematic of the syngeneic subcutaneous flank tumor model. SPF mice were implanted in the flank with B16-F10 tumor cells. Tumor volume measurements began on day 4, when tumors reached approximately 50–100 mm³. Intratumoral (i.t.) injections of diABZI and/or bile acids were administered every three days starting on day 8, when tumors reached ∼250 mm³. (b) B16-F10 melanoma tumor growth in mice treated i.t. with vehicle (DMSO), DCA, diABZI, or the combination (DCA + diABZI). (c) B16-F10 tumor growth in mice treated i.t. with diABZI alone or in combination with CA or DCA. (d-h) Quantification of tumor-infiltrating CD8⁺ T cells expressing Sca-1⁺ (d) CD69 (e) Granzyme B (f) Ki-67 (g), or PD-1 (h) in B16-F10 tumors from mice treated with DMSO, DCA, diABZI, or the combination. Tumors were harvested 2 days after a single i.t. injection. Data in b and c represent mean ± SEM and were analyzed using a mixed-effects model with Tukey’s multiple comparisons post hoc test (n = 8 mice per group). Arrowheads indicate the days of i.t. injection. Data in d**–** h were pooled from two independent experiments, represent mean ± SEM, and were analyzed using an unpaired *t*-test (n = 9–12 mice per group). Each symbol represents one mouse. *P < 0.05, **P < 0.01, ***P < 0.001, ****P < 0.0001; ns, not significant.

To investigate whether treatment-associated alterations in gut microbiota composition contributed to the enhanced antitumor activity, we performed 16S rRNA gene sequencing on fecal samples collected from all treatment groups. Alpha diversity analysis showed that Shannon diversity was maintained across treatment groups, with no significant reduction relative to DMSO-treated controls (**Extended Data Fig. 8a**). Beta diversity analysis revealed modest overall community differences between groups (**Extended Data Fig. 8b**). Linear discriminant analysis effect size (LEfSe) revealed that diABZI treatment was not associated with outgrowth of any dominant bacterial taxon: only minor enrichment of *Allobaculum* was detected, while *Desulfovibrio* and *Bifidobacterium* were modestly reduced (**Extended Data Fig. 8c, d**). DCA treatment was associated with increased abundance of *Allobaculum* and *Turicibacter*, consistent with previous reports that bile acid composition influences the abundance of specific *Firmicutes* taxa. The combination treatment group also exhibited microbiota shifts, including enrichment of *Allobaculum*, *Anaeroplasma*, and *Alistipes*, and a reduction in *Anaerotruncus*, relative to DMSO; however, none of these taxa are established mediators of innate antitumor immunity (**Extended Data Fig. 8c, d**). Together, these results suggest that large-scale microbiota restructuring associated with the combination treatment is unlikely to be the primary driver of the enhanced antitumor activity. Instead, these findings support a model in which DCA promotes STING activation *in vivo*.

To further investigate the antitumor immune response, we analyzed the tumor-infiltrating lymphocytes (TILs), tumor-draining lymph nodes (tdLN) and spleen from B16-F10-bearing mice by flow cytometry (**Supplementary Fig. 1**). diABZI treatment, with or without DCA, significantly upregulated the pro-inflammatory type-I interferon marker Sca-1 across all three tissues, while the combination of DCA and diABZI selectively enhanced Sca-1 expression in TILs (**Fig. 3d, Extended Data Fig. 7e, f**). This increase occurred without changes in the absolute number of CD45^+^ leukocytes or CD3^+^ lymphocytes in TILs (**Extended Data Fig. 7g, h**). However, CD8⁺ T cells from the combination treatment group displayed significantly higher levels of activation markers, including CD69 (**Fig. 3e**), granzyme B (**Fig. 3f**), Ki-67 (**Fig. 3g**), and PD-1 (**Fig. 3h**). In addition, both the frequency and granzyme B expression of NK1.1⁺ cells were elevated in the DCA + diABZI group (**Extended Data Fig. 7i, j**). These results support a model in which DCA enhances STING-associated immune responses within the tumor microenvironment.

To gain insight into the synergistic antitumor effect of DCA + diABZI, we carried out single-cell RNA-sequencing (scRNA-seq) analysis of TILs isolated from mice on day 13 post-tumor implantation. This analysis would also potentially reveal which cell populations were responding to the combination treatment, and how this might impact antitumor immunity. After scRNA-seq analysis, we were able to detect 20 immune cell populations present in tumors from all four treatment groups (**Fig. 4a, b**). NK cells were conspicuously over-represented in the diABZI and especially DCA + diABZI groups (**Fig. 4c**). This was confirmed by statistical analysis, which also revealed overrepresentation of cycling CD8+ T cells and a population of macrophages marked by expression of the complement component C1qa in the DCA + diABZI samples relative to diABZI alone (**Extended Data Fig. 9a**). This was at the expense of B cells, γδT cells, and plasmacytoid dendritic cells (pDCs), which were relatively contracted. As assessed by the number of differentially expressed (DE) genes (adjusted p-value < 0.05; log_2_ FC > 0.5) between DMSO and diABZI groups, the cell populations that responded most robustly to diABZI treatment were NK cells, a cluster of macrophages marked by type I IFN signaling (IFN MF) and monocytes (**Fig. 4d**). Among these cell clusters, as well as additional populations that had at least 10 DE genes in response to diABZI, only NK cells had a substantial response to the combinatorial treatment of DCA + diABZI (**Extended Data Fig. 9b**). Though mild, this response appeared synergistic, as a proportion of genes induced by diABZI were further induced by DCA + diABZI. Scoring each cell for expression of a cell-type specific “diABZI response”, or the aggregate expression of genes induced by diABZI in that cell-type, revealed that NK cells uniquely expressed an enhanced diABZI response in the DCA + diABZI conditions (**Fig. 4e and Extended Data Fig. 9c**). Our analysis additionally revealed that DCA alone did not induce any measurable STING activation across cells responsive to diABZI and that in IFN MFs and monocytes, DCA + diABZI co-administration actually reduced the transcriptional diABZI response (**Extended Data Fig. 9c**). Given that NK cells express components of the STING signaling pathway, but not any canonical receptors for DCA (**Fig. 4f**), this suggested that the synergistic transcriptional response in NK cells was due to enhanced STING signaling by DCA.

**Figure 4.**
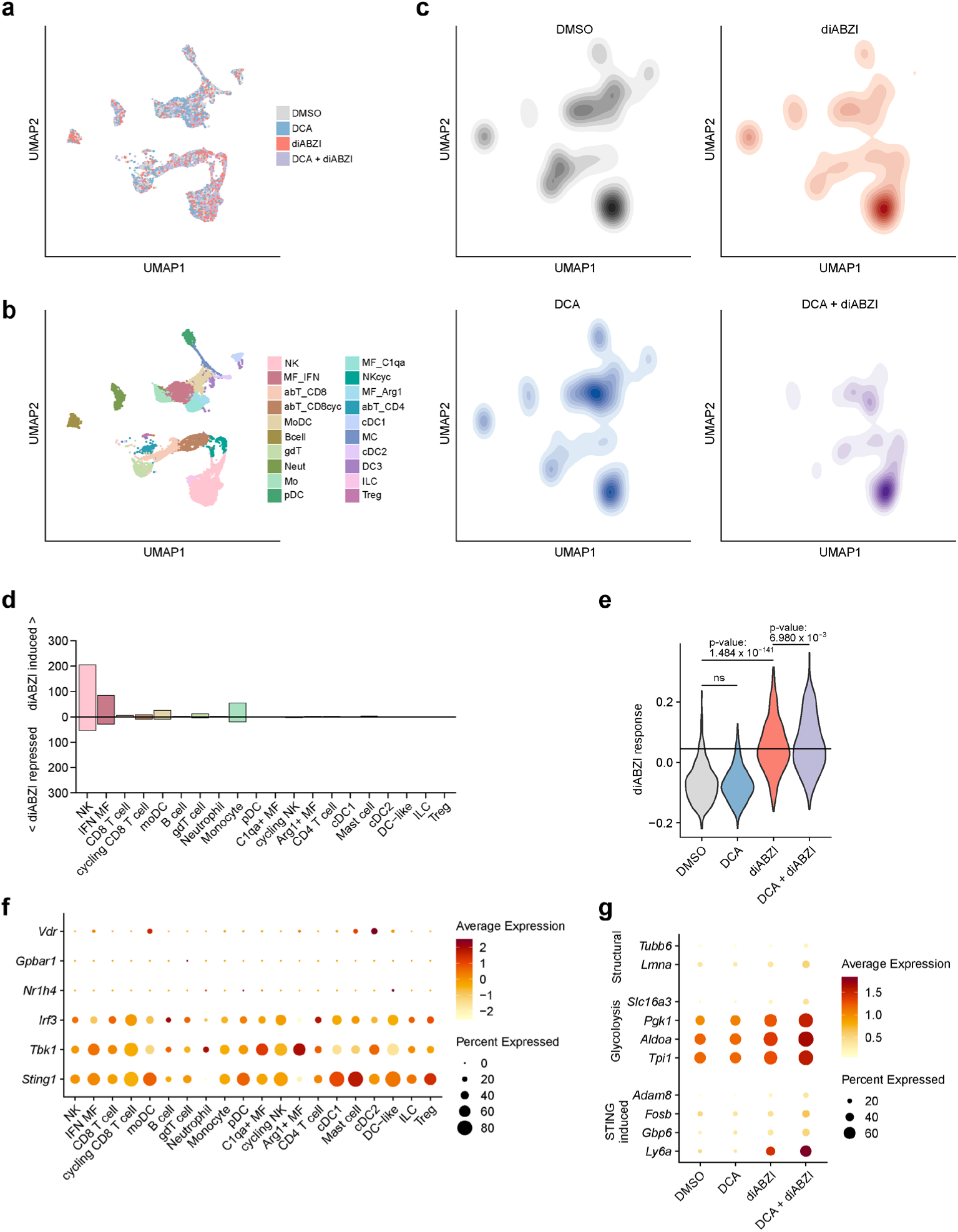
DCA selectively enhances diABZI-induced STING-associated transcriptional responses in tumor-infiltrating NK cells. CD45^+^ immune cells were isolated from tumors of mice (n = 4 per group) subjected to the indicated treatments and analyzed by scRNAseq. UMAP plots with individual cells colored by treatment (a), manually annotated cell type (b), or transformed into a 2D density distribution by treatment group (c). Bar plots showing number of differentially expressed genes in each cell type between DMSO and diABZI groups (d). Cells from each cell type were scored for expression of genes induced by diABZI treatment in that cell type (adjusted p-value < 0.05; log_2_ FC > 0.5). Violin plots (e) show per cell “diABZI” score for NK cells grouped by treatment. P-values determined by ks test. Dot plots showing normalized expression (shading) and percent of cells expressing (size) genes associated with bile acid sensing or STING signaling grouped across all treatments by cell type (f). Dot plots showing normalized expression (shading) and percent of cells expressing (size) select genes significantly induced by DCA + diABZI over diABZI alone in NK cells, grouped by treatment (g).

Focusing on specific genes in NK cells, DCA + diABZI co-administration increased expression of STING targets *Ly6a*, *Gbp6*, *Fosb*, and *Adam8* compared to diABZI alone, which further suggested DCA enhancement of STING-signaling (**Fig. 4g**). The DCA + diABZI elevated expression of genes involved in glycolysis (*Slc16a3*, *Pgk1*, *Aldoa*, *Tpi1*) and intracellular structures (*Lmna*, *Tubb6*) also suggested increased activation and cell growth of NK cells (**Fig. 4g**). These transcriptional indicators of enhanced NK cell activation were also consistent with our flow cytometric analysis. Collectively, these data indicate that DCA enhances STING signaling in the tumor microenvironment specifically in NK cells, potentially driving increased antitumor responses by these cells, although our analysis did not reveal what unique features of NK cells made them especially sensitive to DCA + diABZI treatment relative to other diABZI-responsive immune cell populations. Nonetheless, the results of these analyses were consistent with the notion that DCA potentiated antitumor immunity by enhancing diABZI activation of STING signaling, rather than through some STING-independent activity. Although DCA alone had effects on some immune cells and induced minor gene expression changes (**Extended Data Fig. 9a**), canonical receptors for DCA were poorly expressed across these different immune cell populations (**Fig. 4f**), and the combinatorial treatment specifically enhanced STING-signaling in NK cells, a cell-type important for diABZI-mediated antitumor immunity(*72*).

### Microbiota-producing DCA promote STING activation

Our discovery that DCA modulates STING activation suggests that specific microbiota species capable of producing secondary bile acids may regulate this immune signaling pathway *in vivo*. Of note, specific microbiota species such as *Akkermansia muciniphila* have been reported to secrete cyclic dinucleotides (CDNs) that directly activate the STING pathway and regulate intratumoral IFN-I signaling(*61*). Other microbiota species have also been shown to enhance the efficacy of immune checkpoint blockade (ICB) via STING activation by unknown mechanisms of action(*62–64*). However, whether microbiota-derived secondary bile acids such as DCA can boost the therapeutic efficacy of STING agonists was unknown. To address this, we evaluated STING-mediated IFN-β induction following intraperitoneal injection of diABZI in mice treated with or without a broad-spectrum antibiotic (Abx) cocktail (**Extended Data Fig. 10a**). In specific-pathogen-free (SPF) mice, diABZI induced robust IFN-β expression, whereas antibiotic-treated mice showed markedly decreased IFN-β levels in serum (**Extended Data Fig. 10b**) and cecum (**Extended Data Fig. 10c**). To test whether bile acids can rescue STING activation in antibiotic-treated mice, we co-injected diABZI with bile acids into the Abx-treated mouse (**Extended Data Fig. 10d**) and found that DCA, but not CA, synergized with diABZI to increase serum IFN-β (**Extended Data Fig. 10e**) and IL-6 (**Extended Data Fig. 10f**) levels. In addition, DCA synergized with CDN STING agonists such as c-di-AMP and enhanced serum levels of IFN-β (**Extended Data Fig. 10g**). We also examined the effect of vancomycin (**Extended Data Fig. 10h**), an antibiotic that targets Gram-positive bacteria and has been reported to increase primary bile acids while depleting secondary bile acids in the gut(*18*). Compared to untreated SPF mice, vancomycin-treated mice exhibited reduced levels of DCA in both feces (**Extended Data Fig. 10i**) and serum (**Extended Data Fig. 10j**) as well as reduced diABZI-induction of IFN-β (**Extended Data Fig. 10k**) and IL-6 (**Extended Data Fig. 10l**), demonstrating antibiotic depletion of microbiota and secondary bile acids impaired STING activation *in viv*o.

As vancomycin has been reported to deplete secondary bile acid-producing microbiota species such as *Clostridium scindens*(*18*), we evaluated whether *C. scindens* impacts STING activation *in vivo* compared to *Clostridium sporogenes*, which does not contain the *bai* operon responsible for producing secondary bile acids(*6, 39*). To investigate whether DCA-producing microbiota species modulate STING activation *in vivo*, we colonized antibiotic-treated mice with *C. scindens* or *C. sporogenes* and evaluated their effects on type I IFN activation and antitumor activity. We found that *C. scindens*, which was able to convert CA to DCA *in vitro* (**Extended Data Fig. 10m**), but not *C. sporogenes*, promoted IFN-β induction in vancomycin-treated mice stimulated with STING agonist diABZI (**Extended Data Fig. 10n**). To examine whether *C. scindens* enhances the antitumor efficacy of STING agonists, we colonized antibiotic cocktail or vancomycin-treated mice with bacteria and administered intratumoral injections of diABZI in the B16-F10 tumor model (**Fig. 5a**). Mice colonized with *C. scindens*, but not *C. sporogenes*, showed significantly reduced tumor growth following diABZI treatment in either antibiotic cocktail (**Fig. 5b, Supplementary Fig. 2a**) or vancomycin **(Fig. 5c, Supplementary Fig. 2b**) treated mice. Notably, *C. scindens*, but not *C. sporogenes*, also promoted the antitumor activity of a single diABZI injection and was inactive alone (**Fig. 5d**). Flow cytometry analysis of TILs **(Supplementary Fig. 2c-e**) revealed that *C. scindens*, but not *C. sporogenes* supplementation significantly increased NK1.1⁺CD3⁻ cells (**Fig. 5e, f, Supplementary Fig. 2c-e**), CD3⁺CD45^+^ cells (**Fig. 5g**), and activated Sca-1^+^CD8^+^ T cells (**Fig. 5h**), Ki67^+^CD8^+^ T cells (**Fig. 5i**), and PD-1^+^CD8^+^ T cells (**Fig. 5j**), similar to mice co-injected with DCA and diABZI. Together, these findings demonstrate that DCA-producing microbiota such as *C. scindens* can synergize with STING agonists to enhance type I interferon signaling and antitumor immunity *in vivo*.

**Figure 5.**
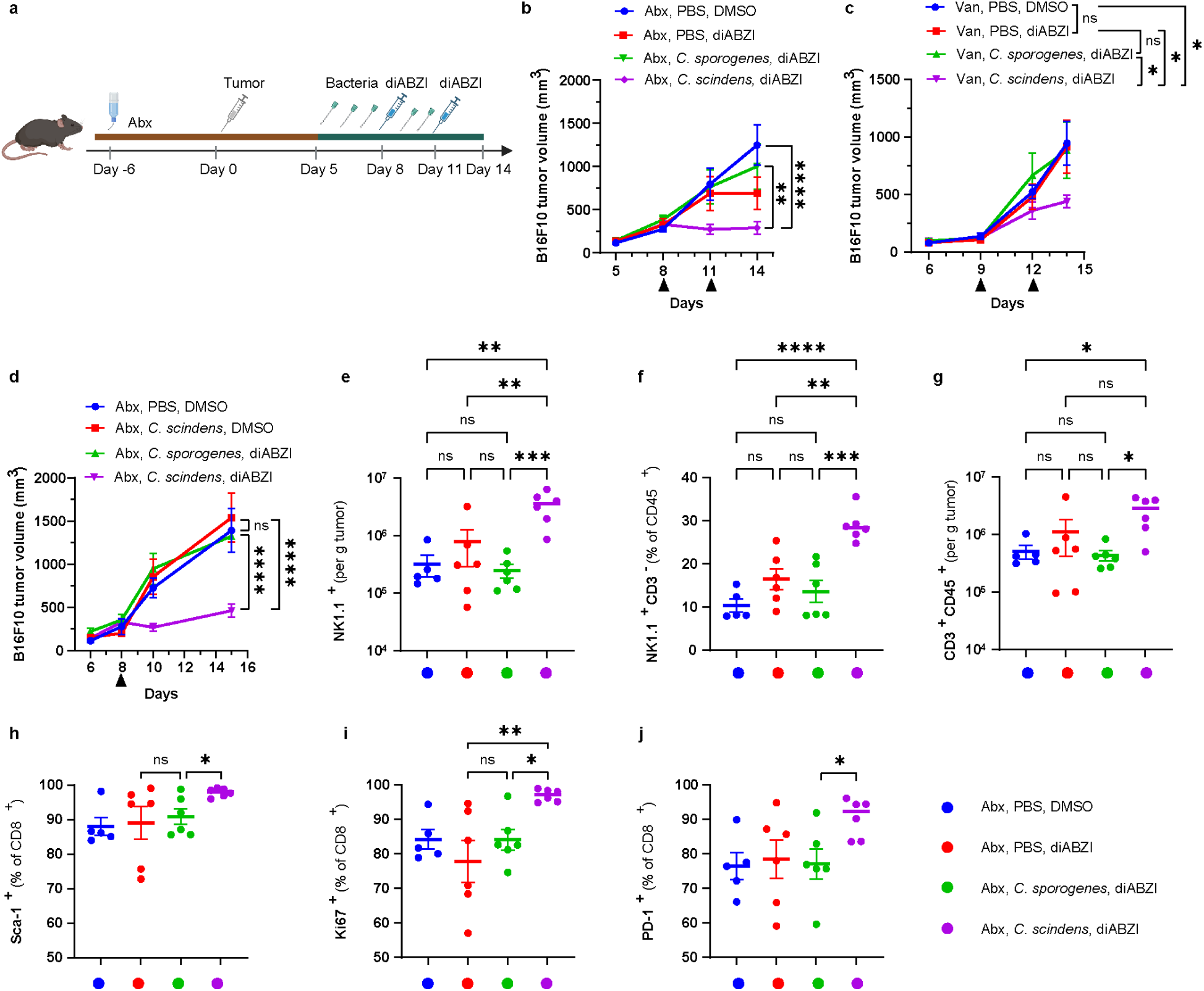
Microbiota-producing DCA promote STING-mediated antitumor responses. (a) Schematic of the tumor growth model used for panels b and c. (b–c) B16-F10 tumor growth curves in mice treated with Abx, n=8 (b) or vancomycin, n=5 (c). Mice received Abx or vancomycin for 10 days. On day 5 post-tumor implantation, Abx or vancomycin treatment was stopped, and mice began daily oral gavage with PBS, *C. sporogenes*, or *C. scindens*. diABZI was administered via i.t. injection on days 8 and 11. (d**-**j) B16-F10 tumor growth (d) and flow cytometry analysis (e-j) in Abx-treated mice gavaged with the indicated bacteria and receiving a single diABZI injection. *n* = 5–6. Absolute number (e) and frequency (f) of tumor-infiltrating NK1.1^+^ CD3^-^ cells, and absolute number of tumor-infiltrating CD3^+^ CD45^+^ cells (g). Frequency of tumor-infiltrating CD8⁺ T cells expressing Sca-1 (h), Ki67 (i), or PD-1 (j). Data in b**-**d are represented as mean ± SEM and were analyzed using a mixed-effects model with Tukey’s multiple comparisons post hoc test. Data for e**-**j represent mean ± SEM and were analyzed using an unpaired two-tailed *t*-test. *P < 0.05, **P < 0.01, ***P < 0.001, ****P < 0.0001; ns, not significant.

To understand whether the antitumor effects of *C. scindens* were associated with enhanced diABZI activity, as well as to compare the effects of bacterially produced DCA to intratumoral administration (**Fig. 4**), we carried out additional scRNA-seq analysis of TILs isolated from tumors of mice treated with vehicle or diABZI, as well as those administered *C. scindens* or *C. sporogenes* in addition to diABZI treatment. After scRNA-seq analysis, we were able to identify the same 20 immune cell populations as in the previous analysis present in tumors from all four treatment groups (**Fig. 4a, b and 6a, b**). Consistent with *C. scindens* administration mimicking DCA injection, NK cells were again conspicuously over-represented with diABZI treatment and *C. scindens* administration (**Fig. 6c**). This was confirmed by statistical analysis, which also revealed overrepresentation of cycling NK cells (**Extended Data Fig. 9d**). As in the DCA experiment, C1qa^+^ macrophages were also proportionally expanded in the *C. scindens* samples relative to *C. sporogenes*, while monocyte-derived DCs (moDCs) were uniquely expanded in this bacterial administration experiment (**Extended Data Fig. 9d**). This was at the expense of B cells, ILCs, Treg cells, and pDCs, which were relatively contracted. We re-defined diABZI induced genes in each cell type for this specific experimental setup and saw that the cell populations that responded most robustly to diABZI treatment were NK cells, a cluster of macrophages marked by type I IFN signaling (IFN MF), and monocytes (**Fig. 4**). Unique to this experiment, CD8^+^ T cells also showed robust induction of gene expression in response to diABZI, consistent with flow cytometric analysis showing stronger CD8^+^ T cell responses in this model (**Fig. 6d**). This unique CD8^+^ T cells responsiveness to diABZI might be due to the antibiotic pretreatment, bacterial administration and/or different tumor growth kinetics in this experiment.

**Figure 6.**
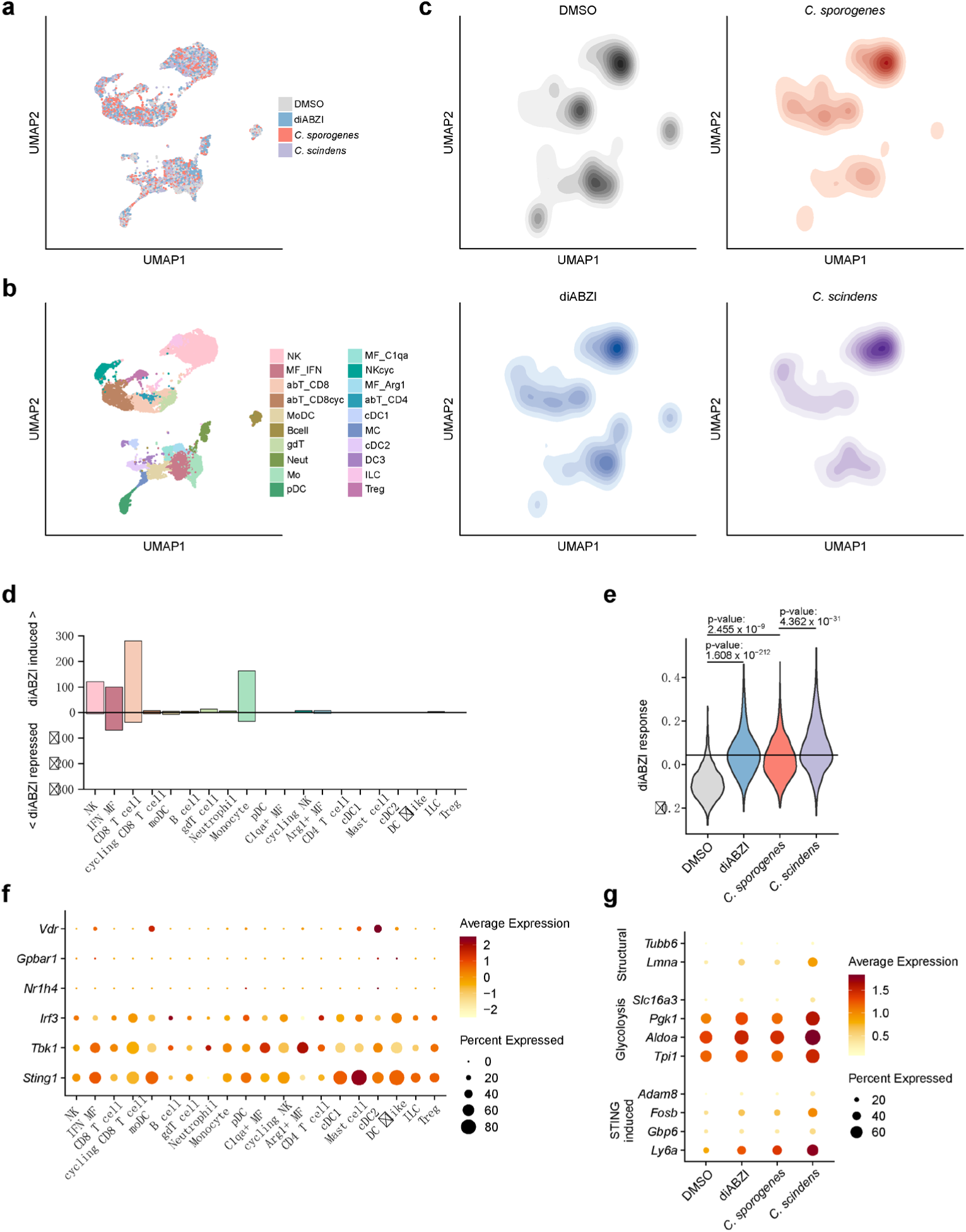
The DCA-producing bacterium *Clostridium scindens* selectively enhances diABZI-induced STING-associated transcriptional responses in tumor-infiltrating NK cells. CD45^+^ immune cells were isolated from tumors of mice (n = 4 per group) subjected to indicated treatments and analyzed by scRNAseq. UMAP plots with individual cells colored by treatment (a), manually annotated cell type (b), or transformed into a 2D density distribution by treatment group (c). Bar plots showing number of differentially expressed genes in each cell type between DMSO and diABZI groups (d). Cells from each cell type were scored for expression of genes induced by diABZI treatment in that cell type (adjusted p-value < 0.05; log_2_ FC > 0.5). Violin plots (e) show per cell “diABZI” score for NK cells grouped by treatment. P-values determined by ks test. Dot plots showing normalized expression (shading) and percent of cells expressing (size) genes associated with bile acid sensing or STING signaling grouped across all treatments by cell type (f). Dot plots showing normalized expression (shading) and percent of cells expressing (size) select genes significantly induced by *C. scindens* (+ diABZI) over *C. sporogenes* (+ diABZI) in NK cells, grouped by treatment (g).

DCA-producing bacteria had strikingly similar effects on diABZI responses as DCA injection, since among these cell clusters, as well as additional populations that had at least 10 DE genes in response to diABZI, only NK cells had a substantial synergistic response to the treatment of diABZI and *C. scindens* relative to diABZI with *C. sporogenes* (**Extended Data Fig. 9e**). For other diABZI-responsive populations such as monocytes and IFN+ MFs, *C. scindens* repressed the diABZI response (**Extended Data Fig. 9e, f**). This is consistent with bacterially derived DCA mimicking the effect of intratumoral injection for these same cell types (**Extended Data Fig. 9b, c**). Scoring each cell for expression of a cell-type specific “diABZI response”, or the aggregate expression of genes induced by diABZI in that cell-type, revealed that NK cells uniquely expressed an enhanced diABZI response in the *C. scindens* group (**Fig. 4e, 6e and Extended Data Fig. 9f**). We again confirmed that NK cells in this model express components to mediate STING signaling (**Fig. 6f**), and again conclude that the synergistic transcriptional response in NK cells was indeed due to enhanced STING signaling by DCA. Focusing on specific genes induced by *C. scindens* treatment in NK cells compared to *C. sporogenes*, increased expression of STING targets *Ly6a*, *Gbp6*, *Fosb*, and *Adam8* further recapitulated DCA enhancement of STING-signaling, while increased expression of genes involved in glycolysis (*Slc16a3*, *Pgk1*, *Aldoa*, *Tpi1*) and intracellular structures (*Lmna*, *Tubb6*) suggested increased activation and cell growth of NK cells, potentially indicating enhanced antitumor immune activity by these cells (**Fig. 4g and 6g**). Given the similarities in the effects of DCA administration and DCA-producing bacteria (*C. scindens*) on NK cell transcriptional responses to the STING agonist diABZI (Figs. 4 and 6), we conclude that intratumoral and microbiota-derived DCA from the gut microbiota can enhance STING agonist signaling and antitumor activity *in vivo*.

## Discussion

The chemical diversity of microbiota-derived metabolites represents a major challenge for defining how the microbiota influences host immunity, disease progression and therapeutic responses(*1, 4, 9*). While previous studies have shown that the microbiota and secondary bile acids can modulate type I interferon signaling(*14*), the molecular mechanism by which DCA enhances type I interferon responses remained unclear. Using a reverse chemical microbiology approach(*73*), our proteomic analysis of microbiota-derived bile acid protein targets revealed an unexpected functional interaction between DCA and the innate immune adaptor STING. Through cell signaling, cellular photo-crosslinking, mutagenesis and cryo-EM studies, we found that DCA, but not its primary bile acid precursor CA, engages the transmembrane dimer– dimer interface of STING and enhances agonist-induced STING oligomeric species and downstream immune signaling. Furthermore, our studies show that DCA and DCA-producing microbiota species can promote type I interferon signaling and antitumor activity of STING agonists *in vivo*. These findings suggest that secondary bile acids can act not only through canonical bile acid receptors, such as FXR and TGR5(*12, 13, 23–25*), or nuclear receptors in immune cells(*26–30*), but also through direct modulation of membrane-associated innate immune signaling proteins.

Our results place DCA within an emerging framework in which STING activity is regulated by multiple ligand-binding sites and membrane-proximal regulatory mechanisms. Canonical cyclic dinucleotides and synthetic CDN mimetics activate STING through its cytosolic ligand-binding domain(*29, 41, 42, 44–52*), whereas small molecules such as C53 and NVS-STG2 engage the STING transmembrane domain and promote higher-order oligomerization and immune activation(*53, 54*). Recent studies further indicate that STING activation is influenced by membrane lipids, including cholesterol and phosphoinositides(*56–60*).

In particular, PtdIns(3,5)P2 has been identified as an endogenous lipid ligand that cooperates with cGAMP to promote STING activation(*58, 59*), and PI4P together with the chemical agonist GNE-6468 was reported to activate STING by inducing transmembrane helix rearrangement(*60*). In this context, DCA may represent a microbiota-derived sterol-like modulator of a lipid- and small-molecule-sensitive STING transmembrane regulatory interface. Whether DCA competes with, cooperates with, or functionally substitutes for endogenous lipid regulators at this interface remains unknown. This mode of action may explain why DCA broadly enhanced responses to structurally distinct STING agonists, including CDNs and non-CDN synthetic agonists, while showing little activity on its own. Thus, DCA may lower the threshold for agonist-induced STING activation by biasing or stabilizing STING conformations that are competent for downstream signaling. Recent systematic mutational profiling further demonstrated that single-amino-acid substitutions throughout STING can broadly tune ligand sensitivity and differentially regulate downstream signaling outputs, highlighting the substantial conformational and functional plasticity of STING(*74*).

These findings also provide a mechanistic link between bile acid metabolism and type I interferon-dependent antitumor immunity. Previous studies have shown that microbiota-derived secondary bile acids can exert both beneficial and pathogenic effects depending on tissue context, concentration and duration of exposure. DCA and related secondary bile acids have been associated with colonization resistance, antiviral immunity and immune modulation(*6, 14–16*), but chronic or elevated exposure has also been linked to tissue injury, inflammation and tumor-promoting immune regulation in liver and colorectal cancer models(*18–21*). Our study reconciles part of this apparent complexity by showing that DCA does not simply act as a general immune stimulant; rather, it can amplify STING agonist-induced signaling in specific therapeutic contexts. In tumor models, DCA or DCA-producing *C. scindens* enhanced the antitumor activity of low-dose diABZI, whereas DCA or *C. scindens* alone had little activity. The differential responses observed across B16-F10, Panc02 and MC38 tumors further suggest that the impact of DCA–STING synergy depends on baseline tumor immunogenicity and responsiveness to STING agonism.

The *in vivo* immune analyses support a model in which DCA and DCA-producing microbiota enhance STING-associated immune activation within the tumor microenvironment. Flow cytometry showed increased activation of tumor-infiltrating CD8+ T cells after DCA plus diABZI treatment and increased NK and CD8+ T cell activation after *C. scindens* plus diABZI treatment. Single-cell RNA-seq further revealed enhanced STING and type I interferon-associated transcriptional programs, with particularly prominent activation signatures in NK cell populations. These findings suggest that the antitumor effects of DCA– STING synergy are not explained solely by changes in total immune cell abundance, but instead involve qualitative activation of innate and adaptive antitumor immune programs. In addition, 16S rRNA gene sequencing did not support selective outgrowth of a dominant immunostimulatory bacterial taxon after diABZI or DCA plus diABZI treatment, arguing that the enhanced antitumor response is unlikely to be driven primarily by broad microbiota restructuring under the conditions tested.

Therapeutically, our findings raise the possibility that bile acid composition and the abundance of DCA-producing microbiota may influence responses to STING-targeted therapies. Because STING agonists can produce dose-limiting inflammatory toxicities and cell type-dependent effects(*53, 54*), strategies that enhance STING agonist activity at lower doses could be valuable. However, the translational implications of DCA–STING synergy require caution. DCA has well-established context-dependent activities, including membrane-active properties at high local concentrations and reported associations with inflammation and tumor promotion in some settings (*18–21, 40*). Therefore, DCA itself may not be an optimal therapeutic agent for systemic modulation of STING. Instead, this work suggests that bile acid-inspired molecules, microbiota-informed patient stratification, or controlled local modulation of secondary bile acid metabolism could be explored as strategies to improve STING agonist efficacy while minimizing risk.

Several limitations should be noted. First, although the cryo-EM density, docking model, photoaffinity competition and mutagenesis data support DCA engagement at or near the STING transmembrane dimer– dimer interface, they do not unambiguously define a single DCA binding pose or exclude dynamic binding modes within this pocket. Additional structural studies with optimized DCA analogues, higher ligand occupancy, or orthogonal biophysical approaches will be needed to fully resolve the binding mode. Second, our bile acid structure–activity analysis suggests that hydroxylation pattern, 3-hydroxyl orientation, steroid ring configuration and side-chain positioning influence DCA-mediated STING modulation, but a more systematic SAR study will be required to define the chemical features that control potency, selectivity and safety. Third, naturally occurring STING alleles and cell-type-specific STING expression may influence the magnitude of DCA-mediated enhancement(*65*), and future studies using human primary cells from genetically diverse donors and cell type-specific *Sting*-deficient mouse models will be important. Germline *Sting*-deficiency would be expected to eliminate the activity of diABZI itself and therefore may not distinguish the specific mechanism by which DCA enhances agonist responses; nevertheless, refined genetic models could help identify the host cellular compartments responsible for DCA–STING synergy *in vivo*.

Finally, our study focused primarily on STING agonist-induced type I interferon signaling and antitumor immunity. Recent work has shown that STING can also function as a proton channel and regulate noncanonical autophagy, lysosomal biology, cell death and inflammasome-related outputs(*75, 76*). Notably, STING proton-channel activity can be mechanistically decoupled from canonical TBK1–IRF3-dependent interferon induction, as C53 blocks STING-dependent proton flux, LC3B lipidation and inflammasome activation while preserving interferon-inducing activity under specific experimental conditions(*75, 76*). We did not systematically determine whether DCA affects STING proton-channel activity, STING-dependent autophagy or inflammasome activation. Under the conditions tested here, DCA mainly enhanced STING-associated type I interferon programs rather than broadly activating other innate immune pathways, but additional studies will be needed to define whether DCA or DCA-inspired molecules differentially tune canonical interferon signaling versus noncanonical STING functions. Moreover, the intratumoral low-dose diABZI regimen used in our mouse tumor models was selected to provide an experimental window for detecting synergy and cannot be directly compared with clinically achievable exposure, particularly because diABZI remains a preclinical agonist and pharmacokinetic parameters vary by compound, dose and route of administration. Future pharmacokinetic, safety and efficacy studies will be required to determine whether modulation of the DCA–STING axis can be therapeutically leveraged.

## Methods

### Cell culture

HEK293T, HT-29, HeLa, THP-1, RAW264.7, B16-F10, MC38, Panc02 were obtained from ATCC. PBMCs were freshly isolated from donor. HEK293T, HT-29, HeLa, RAW264.7, B16-F10, MC38, Panc02 cells were cultured in DMEM (Gibco) supplemented with 10% FBS (GE Healthcare HyClone) and 100 μg ml^−1^ penicillin–streptomycin (Thermo Fisher Scientific). THP-1 and hPBMCs were cultured in RPMI supplemented with 10% FBS. All cells were cultured in a 37 °C incubator with 5% CO_2_. Cells were also routinely cultured without antibiotics to ensure no bacterial infection.

### Bacterial culture

*C. scindens* ATCC 35704 and *C. sporogenes* ATCC 15579 were obtained from C.-J. Guo lab (Weill Cornell Medical School). Bacteria were cultured in Modified Reinforced Clostridial broth (ATCC Medium 2107) and/or TSAB (Tryptic Soy Agar supplemented with 5% Horse Blood, 1% L-Cysteine hydrochloride, and 0.1% Vitamin K3) within an anaerobic chamber (Coy Laboratory Products) at 37 °C for 24 to 48 hours(*77*). Mice were administered 1 × 10^8^ CFU of bacteria daily via oral gavage.

### Animals

Specific pathogen-free, seven-week-old male and female C57BL/6 (B6, 000664) mice were obtained from the Scripps Rodent Breeding Colony. The mice were housed in autoclaved cages with SaniChip bedding and nest-building enrichment on a 12-hour light/dark cycle. They were provided gamma-irradiated chow (LabDiet 5053) and sterile drinking water ad libitum. All animal care and experimental procedures were conducted in accordance with NIH guidelines and were approved by the Institutional Animal Care and Use Committee at Scripps Research (Protocol AUP-21-095).

### Chemical reagents

Compounds were purchased and used directly without further purification. Catalog is shown as below: diABZI (99.84%, MCE, HY-112921B), MSA-2 (≥98%, Cayman, 30140), c-di-AMP (≥ 95% by LC/MS, InvivoGen, tlrl-nacda), 2’3’-cGAMP (≥ 95% by LC/MS, InvivoGen, tlrl-nacga23), c-di-GMP (≥ 95% by LC/MS, InvivoGen, tlrl-nacdg), C53 (≥98%, Cayman, 37354), cholic acid (97%, Alfa Aesar, A11267), NVS-STG2 (98%, MCE, HY-157214), DMXAA (≥98%, Cayman, 14617), BB-Cl-Amidine (≥95%, MCE, hy-111347), single-stranded template DNA (HT-DNA) (sigma, D7656), deoxycholic acid (≥98%, HPLC, Sigma, D2510-100G), chenodeoxycholic acid (>98%, Abcam, ab142267), lithocholic acid (≥95%, Sigma, L6250-25G), ursodeoxycholic aicd (99%, BTC, 123710-5g), Alloisolithocholic Acid (≥95%, Cayman, 29542), TCA (≥97%, sigma, 86339-1G), GCA (≥95%, SC-218574A), TDCA (SC-253596), GDCA (≥97%, HPLC, sigma, G9910-1G), TCDCA (99%, spectrum, T3925), GCDCA (spectrum, G3389), PEG400 (MCE, HY-Y0873A), Az-biotin was purchased from Sigma-Aldrich (762024).

### Photoaffinity probe cell treatments

THP-1 cells were maintained in RPMI 1640 medium supplemented with 10% fetal bovine serum (FBS), ensuring a cell density below 1 × 10^6 cells/mL. To initiate treatment, cells were harvested by centrifugation at 200 g for 10 minutes, the supernatant was removed, and the pellet was resuspended in serum-free RPMI 1640 containing the bile acid reporter (CA2 -25 µM, DCA2-25 µM, DCA4-25 µM). The cells were then incubated for 30 minutes at 37 °C in a humidified atmosphere of 5% CO₂. Following incubation, the medium was removed and replaced with cold DPBS, and the cells were placed in 15 cm dish on ice and exposed to UV light (365 nm) for 10 minutes and transferred into tubes. The suspensions were centrifuged again at 200 g for 10 minutes, and aspirated to remove the supernatant. Finally, the resulting cell pellets were flash-frozen in liquid nitrogen and stored at –80 °C until further processing.

### Preparation of samples for chemoproteomic analysis

Chemoproteomic samples were prepared in biological duplicates for each condition using previously described procedures(*78*). Briefly, cell pellets were resuspended in 500 µl of cold DPBS and lysed via sonication (Branson Sonifier probe; 15 ms on, 40 ms off, 15% amplitude, 7 pulses). Protein concentrations were normalized to 2 mg/ml in 500 µl of ice-cold DPBS, using a DC Protein Assay (Bio-Rad), in 15-ml falcon tubes. The “click-chemistry cocktail” was then added to each lysate, consisting of tris[(1-benzyl-1H-1,2,3-triazol-4-yl)methyl]amine (TBTA, final concentration: 100 µM), tris(2-carboxyethyl)phosphine (TCEP, final concentration: 1 mM), biotin-azide (final concentration: 100 µM), and CuSO_4_ (final concentration: 1 mM). Samples were incubated at room temperature for 1 hour. Ice-cold 4:1 MeOH/CHCl_3_ (2.5 ml) and ice-cold DPBS (1 ml) were added to each sample to quench the click reaction, followed by centrifugation (3,200 g, 4 °C, 10 min) to precipitate proteins. The precipitates were washed twice more with 4:1 MeOH/CHCl_3_ and resuspended in 500 µl of 6 M urea in DPBS, along with 10 µl of 10% SDS. Protein reduction was performed by adding 50 µl of a freshly prepared 1:1 solution of TCEP (25 µl, 200 mM in DPBS) and K_2_CO_3_ (25 µl, 600 mM in DPBS), and incubating at 37 °C for 30 minutes. Following reduction, proteins were alkylated by adding 70 µl of freshly prepared iodoacetamide (400 mM in DPBS) and incubated for 30 minutes at room temperature in dark. After alkylation, 130 µl of 10% SDS was added to the samples, followed by dilution with 5.5 ml of DPBS. A 100 µl aliquot of 50% streptavidin agarose slurry (Thermo Fisher Scientific, 20349) was added to each tube, and samples were rotated for approximately 1.5 hours at room temperature to enrich probe-labeled proteins. The beads were then centrifuged down to tube bottom (750 *g*, 2 min, 4 °C) and washed once with 0.2% SDS in DPBS (5 min), twice with DPBS, once with water, and transferred to LoBind microcentrifuge tubes with 100 mM TEAB (pH 8.5).

Bound proteins were digested overnight with trypsin (Promega, V5111) in digestion buffer [100 µl, 100 mM TEAB (pH 8.5), 100 µM CaCl_2_] at 37 °C. The resulting peptides were transferred to new tubes for each condition and labeled with the respective tandem mass tags (TMT; Thermo Fisher Scientific) 10plex reagents (8 µl, 20 µg/µl) for 1 h, quenched by hydroxylamine (6 µl, 5% *v*/*v*) for additional 15 min, acidified by formic acid (8 µl), and dried through SpeedVac vacuum concentrator. The dried peptides were redissolved in 0.1% TFA in H_2_O, combined, and proceeded to be fractionated into 12 fractions by Pierce high pH Reversed-Phase Fractionation Kit (Thermo Fisher Scientific, 84868) according to manufacturer’s instructions. The fractions were pooled pairwise into 6 final fractions, dried, and stored at −80°C until mass spectrometry analysis.

### LC-MS analysis of chemoproteomic samples

Proteomic samples were resuspended in MS sample buffer (20 µl, 5% acetonitrile, 0.1 % formic acid in water) prior to LC-MS analysis, following published methods(*78*). In short, 3 µl of each sample was loaded onto an Acclaim PepMap 100 precolumn (75 µm × 2 mm) and separated on an Acclaim PepMap RSLC analytical column (75 µm × 15 cm) using an UltiMate 3000 RSLCnano system (Thermo Fisher Scientific). Buffer A (0.1% formic acid in H_2_O) and buffer B (0.1% formic acid in acetonitrile) were used in a 220-minute gradient at a flow rate of 300 nl/min. The gradient consisted of 2% buffer B for the first 10 minutes, a gradual increase to 30% buffer B over the next 192 minutes, 60% buffer B for 5 minutes, followed by a ramp to 95% buffer B for 1 minute, held for 5 minutes, then lowered back to 2% buffer B for 1 minute, and re-equilibrated for 6 minutes.

Eluents were analyzed using a Thermo Fisher Scientific Orbitrap Fusion Lumos mass spectrometer with a 3-second cycle time and a nano-LC electrospray ionization voltage of 2,000 V. Data were acquired with Xcalibur software (v4.1.50). MS1 spectra were captured with a scan range of 375–1,500 m/z, a maximum injection time of 50 milliseconds, and dynamic exclusion enabled (1 repeat, exclusion duration of 20 seconds). The resolution was set to 120,000 with an AGC target of 1 × 10^6^. Peptides chosen for MS2 analysis were isolated with a quadrupole (isolation window of 1.6 m/z) and fragmented by collision-induced dissociation (CID) at 30% collision energy. Fragment ions were recorded in the ion trap (AGC target 1.8 × 10^4^, maximum injection time of 120 ms). MS3 spectra were generated using high-energy collision-induced dissociation (HCD) mode with 65% collision energy, and synchronous precursor selection (SPS) was set to 10 SPS ions in MS3 for TMT fragmentation.

### Proteomic result analysis

Proteomic analysis was carried out using Proteome Discoverer (v2.4 or v3.0, Thermo Fisher Scientific) following previously established protocols(*78*). Peptide identification was conducted using the SEQUEST HT algorithm, precursor mass tolerance is 10 ppm and fragment mass tolerances is 0.6 Da, allowing for one missed cleavage. Variable modifications included oxidation (M, +15.994915), carbamidomethyl (C, +57.02146) and TMT-tag modifications (K and N-terminal, +229.1629) were used as fixed modifications. The spectral data were searched against the Homo sapiens proteome database (Uniprot, 2018; 42,358 sequences), applying a 1% false discovery rate (FDR) with Percolator. MS3 peptide quantification used a mass tolerance of 20 ppm, and channel abundances were normalized to the median signal intensity across all channels.

To ensure data reliability, proteins reported required at least two unique peptides. TMT ratios calculated in Proteome Discoverer were transformed using log2(x), and statistical significance of the ratios was assessed with two-tailed t-tests across two biological replicates (significance threshold set at P < 0.05). Detailed proteomics data are available in Supplementary DataS1.

### Pull-down of STING by bile acid photoaffinity probes

From the bile acid photoaffinity probes-treated or control total cell lysates prepared as described above, 225 μL of each total cell lysates (∼250 μg protein) was added with 25 μL of click chemistry reagents as a 10X master mix (Az-biotin: 0.1 mM, 10 mM stock solution in DMSO; tris(2-carboxyethyl)phosphine hydrochloride (TCEP): 1 mM, 50 mM freshly prepared stock solution in dH_2_O; tris[(1-benzyl-1H-1,2,3-triazol-4-yl)methyl]amine (TBTA): (0.1 mM, 2 mM stock in 4:1 t-butanol: DMSO); CuSO_4_: 1 mM, 50 mM freshly prepared stock in dH_2_O). Samples were mixed well and incubated at room temperature for 1 h. After incubation, samples were mixed with 1 mL cold methanol and incubated at -20 °C overnight. Sample proteins were precipitated at 18000 g for 15 min at 4 °C. After gently removing the aqueous layer, protein pellets were washed with 500 μL cold methanol three times, spinning down at 18000 g for 15 min at 4 °C, and liquid was gently decanted. After last wash, pellets were let air dried before being re-solubilized in 100 uL 4% SDS PBS with bath sonication. Solutions were then diluted with 300 µL PBS, spun down (18000 g, 10 min) to remove any particles. 20 µL of supernatant was kept for “input” before enrichment. The left supernatant was incubated with 20 µL PBS-T-washed High Capacity NeutrAvidin agarose (Pierce) (500 µL PBS-T-washed twice, 2500 g, 60 s) at room temperature for 1 h with end-to-end rotation. The agarose was washed with 500 µL of 1% SDS PBS 3 times, 500 µL of 1M Urea PBS 3 times, and 500 µL of PBS 3 times. Samples were boiled with 2X Laemmli buffer 95 °C for 5 min before being loaded onto a 4-20% Tris-HCl gel (Bio-Rad) for SDS-PAGE. Equal loading is validated by immunoblotting of input samples before enrichment (“Input”).

### Western Blot

Cells were lysed by lysis buffer (1% SDS, 20 mM HEPES, 150 mM NaCl, pH 7.4, protease inhibitor cocktail (Roche), phosphatase inhibitor cocktail (PhosSTOP, Roche), benzonase (1:1000)). The concentration of cell lysis was determined by BCA assay (Pierce™ BCA Protein Assay Kits) and normalized with lysis buffer. Samples were then boiling with 1× Laemmli buffer at 95 °C for 5 min and loaded onto a 4–20% Tris-HCl gel (Bio-Rad) for SDS-PAGE. Proteins were then transferred onto a 0.45-μm nitrocellulose membrane (Bio-Rad) with a Trans-Blot Turbo Transfer System (Bio-Rad) at 25 V for 30 min. The membrane was blocked with 5% non-fat milk in PBS-T or Protein-Free Blocking Buffer (927-80003, LI-COR) (for detection of phosphorylated proteins) for 60 min. Corresponding antibody was added to the solution before incubating the membrane at 4 °C overnight. The membrane was washed with PBS-T three times and incubated with 1:10,000 anti-rabbit HRP (Abcam, ab97051) or anti-mouse HRP (Abcam, ab97023) in PBS-T with 5% non-fat milk at room temperature for 1 h. The membrane was washed with PBS-T three times and imaged with Clarity Western ECL substrate (Bio-Rad) and a ChemiDoc imaging System (Bio-Rad). Following antibodies were used as per the manufacturer’s instructions. NR1H4 Polyclonal antibody (25055-1-AP, Proteintech); IRF-3 (D83B9) Rabbit mAb (4302, Cell signaling technology); Phospho-IRF-3 (Ser386) (E7J8G) XP® Rabbit mAb (37829, Cell signaling technology); TBK1/NAK (D1B4) Rabbit mAb (3504, Cell signaling technology); Phospho-TBK1/NAK (Ser172) (D52C2) XP® Rabbit mAb (5483, Cell signaling technology); STING (D2P2F) Rabbit mAb (13647, Cell signaling technology); Phospho-STING (Ser366) (E9A9K) Rabbit mAb (50907, Cell signaling technology); Tubulin (ab6046, Abcam).

### RT-qPCR

THP-1 or RAW 264.7 cells were grown in 6-well plates to 90% confluency. Cells were then treated with DMSO, DCA, di-ABZI with indicated concentration for 4 hours. RNA was purified using RNeasy Plus Kit (Qiagen). cDNA was then synthesized from purified RNA with iScript advanced cDNA synthesis kit (BioRad). The cDNA was diluted ten times with nuclease-free water. RT-qPCR was performed with PowerUp SYBR Green Master Mix (Applied Biosystems), per the manufacturer’s instructions. The expression of target genes was normalized to the expression of housekeeping gene ACTB. Following primers were used:

*humanIFNB1*
(5’-AAACTCATGAGCAGTCTGCA-3’+5’-AGGAGATCTTCAGTTTCGGAGG-3’)
*humanIL6*
(5’-ACTCACCTCTTCAGAACGAATTG-3’+5’-CCATCTTTGGAAGGTTCAGGTTG-3’)
*humanCXCL10*
(5’-GGTGAGAAGAGATGTCTGAATCC-3’+5’-GTCCATCCTTGGAAGCACTGCA-3’)
*murineIfnb1*
(5’-TCCGAGCAGAGATCTTCAGGAA-3’+5’-TGCAACCACCACTCATTCTGAG-3’)
*murineIl6*
(5’-AACGATGATGCACTTGCAGA-3’+5’-GAGCATTGGAAATTGGGGTA-3’)
*murineCxcl10*
(5’-CCAAGTGCTGCCGTCATTTTC-3’+ 5’-GGCTCGCAGGGATGATTTCAA-3’)

### Dual Luciferase reporter assay

HEK293T cells were seeded in 96-well Flat Clear Bottom White Polystyrene TC-treated Microplates (Corning) at 30,000 cells per well. After 16 h, cells were transiently transfected with three plasmids, including pMSCV-hygro-STING (Addgene, 102598), IFN-Beta_pGL3 (Addgene, 102597), and pRL-TK (Promega, E2241) using Lipofectamine 3000 Transfection Reagent (Invitrogen, L3000-015). After 24 h, cells were treated with indicated compounds for another 24 h. Luminescence was detected after the addition of dual-luciferase reporter assay reagent (Promega, E1960), using a BioTek Synergy Neo2 Multi-Mode Microplate Reader with a 0.5 s integration time. Firefly luciferase luminescence normalized to the Renilla luciferase control is shown.

### STING-GFP imaging

HeLa cells stably expressing STING-GFP were generated by lentivirus transduction and selected by puromycin and GFP cell sorting. Cells grown on glass coverslips were treated with indicated conditions. Cells were then rinsed with ice-cold 1x PBS for 3 times and fixed with 4% paraformaldehyde for 20 minutes. Post fixation, the cells were further washed with 1x PBS for 5 minutes 3 times. Cells were then permeabilized with 0.3% Triton-X in PBS for 20 minutes and washed again with 1x PBS for 5 minutes 3 times before mounting for imaging. DAPI in the mounting medium (VECTASHIELD, H-1800) is used as the counterstain. All incubations were conducted at room temperature unless otherwise specified. Mounted coverslips were imaged with Nikon C2+ confocal microscopy. Resulting images were analyzed with CellProfiler (pipeline provided).

### STING-oligomerization analysis

THP1 cells were seeded in 6-well plates at a density of 1 × 10⁶ cells/mL and incubated with 100 nM or 200 nM diABZI in the presence or absence of 50 μM DCA for 2 hours. Following treatment, the cells were resuspended in lysis buffer containing 50 mM HEPES (pH 7.4), 150 mM NaCl, 5 mM MgCl₂, 1 mM DTT, protease inhibitor cocktail (Roche), and phosphatase inhibitor cocktail (PhosSTOP, Roche). The cell suspension was incubated on ice for 15 minutes, followed by sonication for cell lysis. Lysates were then centrifuged at 1,000 g for 5 minutes to obtain the S1 supernatant. An equal volume of buffer containing 2% NP-40 was added to the supernatant. Protein samples were mixed with 20% glycerol and resolved using a 4–20% Criterion TGX stain-free gel (Bio-Rad). Samples were run in Tris-Glycine running buffer supplemented with 0.05% Coomassie Brilliant Blue G-250 at a constant voltage of 120 V for 30 minutes. Following the initial run, the buffer was exchanged for clear Tris-Glycine buffer to enhance contrast on the gel, and the separation continued until the dye front reached the bottom. Proteins were then transferred to a nitrocellulose membrane via immunoblotting for further analysis. The samples were also denatured and run with SDS-PAGE for immunoblotting with the indicated antibodies.

### Cryo-EM sample preparation

The gene for human STING (UniProtKB Q86WV6) was codon-optimized, synthesized, and cloned into the pEZT-BM expression vector. The C-terminal tail following residue 343 was replaced with a linker region and Strep-tag (GGGSGGGSGGSAWSHPQFEK). HEK293F cells were grown at 37 °C to a density of ∼1 million cells/mL and transfected with 1 mg DNA and 3 mg PEI per liter of cells. After 72 hours, cells were harvested by centrifugation, resuspended in 20 mM Tris 7.4 + 150 mM NaCl (TBS buffer) with 1 mM PMSF and 0.5 mM TCEP, and lysed by sonication. The cell lysate was centrifuged at 3,000 g for 10 min, and the supernatant was further centrifuged at 186,000 g for 2 hours to isolate membranes. The membrane fraction was then stored at -80°C. Membrane fractions were homogenized and solubilized for 2 hours at 4 °C in TBS + 1% DDM, 0.2% CHS, 1 mM PMSF, 0.5 mM TCEP, and 1 mM CaCl_2_, then subsequently clarified via centrifugation at 186,000 g for 40 min. STING protein was affinity-purified from solubilized membranes with Strep-Tactin XT 4Flow (IBA) resin. Protein was eluted with Buffer BXT (IBA), and eluted protein was subjected to gel-filtration using a Superdex 200 Increase 10/300 GL column (Cytiva) in SEC buffer (TBS + 0.02% DDM, 0.004% CHS, 100 µM DCA, and 0.5 mM TCEP). SEC-purified STING was diluted to ∼5 µM and incubated with 30 µM 2’3’-cGAMP (InvivoGen) and 30 µM C53 (Cayman) at 4 °C overnight, after which the sample was concentrated to ∼8 mg/mL. 3 µL of sample was applied to glow-discharged UltrAuFoil 1.2/1.3 300-mesh grids (Quantifoil) before being blotted and plunge-frozen into liquid ethane using a Vitrobot Mark IV (Thermo Fisher).

### Cryo-EM data collection and processing

A total of 5839 cryo-EM micrographs were collected on a 200 kV Glacios 2 microscope equipped with a Falcon 4i detector (Thermo Fisher) at a nominal magnification of 190,000x resulting in a pixel size of 0.718 Å. Data were collected with EPU (Thermo Fisher) with an approximate dose of 45 e/Å2 and a nominal defocus range of -0.6 to -1.5 μm. Micrographs were aligned using Patch Motion Correction and CTF estimation was done by Patch CTF in Cryosparc Live(*79*). Micrographs with CTF fits lower than 8 Å were discarded, and all further data processing was done with Cryosparc. Blob picker was initially used on a subset of 300 micrographs to generate 2D classes to be used as templates for template picking for the full dataset. Picked particles were subjected to multiple rounds of 2D classification to remove false positives. Heterogeneous refinement, using PDB 7SII low-pass filtered to 20 Å as an initial model, was used to select a final subset of 285k particles. Non-uniform refinement(*80*) and local refinement with a mask around two STING protomers yielded a final map at 2.9 Å resolution using C2 symmetry and 3.0 Å with C1 symmetry. PDB 7SII was docked into the C2 map using UCSF ChimeraX(*81*). The model was manually adjusted using Coot (*82, 83*) and subjected to real-space refinement in Phenix(*84*). Structural figures were made using UCSF ChimeraX.

### Docking Method

The STING tetramer structure was protonated using Reduce(*85*) and prepared for docking using Meeko (https://github.com/forlilab/Meeko) to assign atom types and Gasteiger partial charges and convert to a PDBQT file. Grid maps were initially calculated with AutoGrid4 with a box size of 32 Å, 34 Å, 30 Å (0.375 Å grid spacing) and with the box center at 112 Å, 116Å, 92 Å. Ligand files were made into 3-dimensional SD Files from SMILES using Scrubber (https://github.com/forlilab/scrubber), then converted to PDBQTs with Meeko.

The five different ligands (Cholic acid, DCA, Iso-DCA, Iso-LCA, and UDCA) were docked with STING using a custom docking engine employing a search method similar to AutoDock-Vina (Monte-Carlo global search, L-BFGS local search) with the AutoDock4.2 force field and affinity grid maps modified using our CryoXKit software (manuscript submitted) to include guidance from the C2-symmetric cryo-EM density. For creating the density cloud representations of the ligands during docking, we add a Boltzmann weight, *exp*(−*E_n_/kT*), based on the energy *E_n_* of each accepted pose to a grid map at the locations of each atom. This is done separately for each Monte-Carlo run and then subsequently summed into a single map by renormalizing each run’s map points with the overall best energy’s Boltzmann weight *exp*(−*E_best_/kT*). Subsequently, a Gaussian convolution with a width of (1.58/2) Å is performed and the resulting grid map is output as an MRC file.

### ELISA

Mice were pre-treated with either a broad-spectrum antibiotic cocktail (5 g/L streptomycin, 1 g/L colistin sulfate, and 1 g/L ampicillin) (*68, 69*) or vancomycin (0.5g/L) administered in drinking water for one week prior to treatment. On Day 7, mice were ip. injected with STING agonists and/or bile acids. The following doses were used per injection: CA or DCA – 20 mg/kg; diABZI – 1 mg/kg; and c-di-AMP – 1 or 5 mg/kg. For the CA-bacteria experiment, the antibiotic cocktail was replaced with 1mM cholic acid sodium salt (Thermo Fisher, 361-09-1), administered in drinking water in parallel with daily oral gavage of bacteria. Blood samples were collected via retro-orbital bleeding 4 hours post-treatment, and serum was isolated by centrifugation. IFN-β and IL-6 levels were quantified using a mouse IFN-β ELISA kit (PBL Assay Science, 42400-1) and a mouse IL-6 ELISA kit (Thermos Fisher, KMC0061), following the manufacturer’s instructions.

### LC-MS analysis of extracted bile acids

Protocols were adapted from ref 38. Fecal pellets were collected, weighed and transferred into tubes containing several 1 mm diameter beads (Cat. No. 11079110). Each sample was extracted three times with 400 µL cold LC-MS grade methanol, homogenized at 4 m/s for 30 s, and pooled. The combined supernatant (∼1.2 mL) was incubated at –20 °C for 1–2 h to precipitate proteins, then centrifuged at ≥13,000 × g for 15 min at 4 °C. 30 µL of serum samples were treated with 120 µL of cold methanol. The mixture was vortexed and sonicated for 10 min. Proteins were precipitated at −20 °C for 16 hours. Next, the proteins were centrifuged at 13,000 rpm for 15 min at 4 °C. Similarly, bile acids were extracted from 1 mL of sterile-filtered bacterial supernatants with 4 mL of cold methanol for 16 hours, followed by centrifugation at ≥13,000 × g for 15 min at 4 °C. The clear extracts from fecal pellets, serum or bacterial cultures were further subjected to bile acid analysis by LC-MS.

Samples were analyzed using 1290 Infinity II LC/MSD system (Agilent) equipped with Zobrax SB-C18 Rapid resolution HT (2.1 × 50 mm,1.8 μm). Samples were run at flow rate 0.35 mL/min in mobile phase (A: water, 0.1% formic acid), room temperature and an eluent (B: acetonitrile, 0.1% formic acid) using following gradient: 0-1 min: 5% B, 1-8 min: 60-70% B, 8-9 min: 70-95% B, 9-11 min: 95% B, 11-12 min: 95-5% B. Selected ion abundance was collected with MSD API-ES in negative-ion SIM mode with target masses of CA (m/z = 407.2) and DCA (m/z = 391.2). Concentrations of bile acids (CA and DCA) were quantified by calibration with standards and normalized to weight of fecal pellets, volume of serum or bacterial supernatants. Extracted intensities and quantifications were plotted in GraphPad Prism.

### Tumor inoculation, measurements and treatments

Tumor cells were cultured for one week and split the day prior to harvesting for tumor inoculation. Cells were harvested using TrypLE Express (Invitrogen, 12604021), washed, and resuspended in cold phosphate-buffered saline (PBS), then counted using a hemocytometer. The cells were resuspended at 2× concentration in PBS on ice and subsequently diluted 1:1 with Matrigel Matrix (Corning, 356231). Animals were anesthetized with 2.5% isoflurane, and the right flank was shaved using hair clippers. Each animal was subcutaneously injected with 100 μL of the cell suspension into the mid-right flank. The final number of injected cells per cell type was as follows: B16/F10 – 1 × 10⁵ cells; MC38 – 3 × 10⁵ cells; Panc02 – 3 × 10⁵ cells. For the bacterial experiment, mice were injected with 1 × 10⁶ B16/F10 cells without Matrigel. Once tumors were established (approximately 100 mm³), tumor volume was measured every three days using digital calipers and calculated as: volume = length × width² × 0.5, where the width was the smaller of the two measurements. Treatment with diABZI and/or bile acids began when tumors reached ∼250 mm³ (Day 8) and was administered via intratumoral injection every three days. The doses per injection were as follows: CA or DCA – 2.5 mg/kg; diABZI – 50 µg/kg. Mice were euthanized when the tumor volume exceeded 2000 mm³. For the survival assay, mice were considered dead when their tumor volume exceeded 2000 mm³.

### Cell isolation and flow cytometry analysis

Tumors, spleens, and tumor-draining lymph nodes (tdLNs) were dissected from mice. For tumor processing, excess stromal tissue was removed, and the tumor was placed in RPMI containing 1.5 U/mL Liberase TM (Roche, 5401119001), 0.2 mg/mL DNase I (Worthington Biochemical, LS002006), and ceramic spheres (6.35 mm, MP Biomedicals, 116540424-CF). The tissue was incubated at 37 °C with gentle shaking for 30 minutes, followed by passage through a 70 µm filter. For spleen processing, the spleen was transferred to a strainer, mashed using the end of a syringe, washed with RPMI, and centrifuged. Tumor and spleen samples were resuspended in 1 mL of red blood cell lysis buffer (Thermo Fisher, 00-4333-57) for 1–2 minutes at room temperature. For lymph nodes, tissues were directly mashed using the end of a 3 mL syringe and collected in 1 mL RPMI. Tumor, spleen, and tdLN cell suspensions were passed through 100 μm, 44% open area nylon mesh and transferred to 96-well plates.

Cells were washed and stained with 1:1000 Zombie Yellow viability dye (BioLegend, 423103) for 20 minutes at room temperature in the dark. After two washes, samples were incubated with 20 µL of staining buffer containing 0.5 µL TruStain FcX anti-mouse CD16/32 blocking agent (BioLegend, 101319) for 30 minutes at room temperature. Cells were then stained for 30 minutes on ice in the dark with the following antibodies: anti-CD4 (BUV496, GK1.5, BD Biosciences 612953), anti-CD44 (PerCP-Cy5.5, IM7, BioLegend 103031), anti-Sca-1 (BUV395, D7, BD Biosciences 563990), anti-CD69 (PE, H1.2F3, BioLegend, 104507) and anti-PD-1 (BV711, CD279, BD Biosciences 135231). Following staining, cells were washed twice, fixed, and permeabilized using the FoxP3/Transcription Factor Staining Buffer Set (Thermo Fisher, 00-5523-00) overnight at 4 °C. After three washes with permeabilization buffer, cells were incubated with 20 µL of permeabilization buffer containing 10% rat serum (Thermo Fisher, 24-5555-94) for 20 minutes on ice. Subsequently, cells were stained for 20 minutes on ice with the following antibodies: anti-CD45 (APC-Fire 750, 30-F11, BioLegend 103153), anti-CD3 (BV785, 17A2, BioLegend 100232), anti-NK1.1 (BV480, PK136, BD Biosciences 746265) anti-CD8 (PE-Cy-7, 53-6.7, BioLegend 100721), anti-FoxP3 (AF532, FJK-16s, Thermo Fisher 58-5773-80), anti-Granzyme B (PE-CF594, GB11, BD Biosciences 562462) and anti-Ki67 (FITC, SolA15, LifeTech 11-5698-82). Samples were analyzed using Cytek Aurora spectral flow cytometer, and data were analyzed using FlowJo Version 10.9.0.

### 16S rRNA gene sequencing and microbiome analysis

Fecal pellets were collected from individual mice at the experimental endpoint and stored at −80 °C until processing. Microbial DNA was extracted using the Quick-DNA Fecal/Soil Microbe Miniprep Kit (Zymo Research, D6010) according to the manufacturer’s instructions. Amplicon libraries targeting the bacterial 16S rRNA gene hypervariable regions were prepared and sequenced by Azenta Life Sciences (formerly GENEWIZ). Sequencing was performed on an Illumina MiSeq platform using paired-end 2 × 250 bp chemistry (20 samples total; n = 4 mice per group). Raw reads were quality filtered, merged, and screened for chimeric sequences. High-quality reads were clustered into operational taxonomic units (OTUs) at 97% sequence identity using VSEARCH and assigned taxonomic classifications using the SILVA database. Samples were rarefied to an even sequencing depth prior to diversity analyses. Good’s coverage exceeded 99.9% for all samples, yielding 443 OTUs across the dataset. Alpha diversity was assessed using the Shannon diversity index and compared among groups using the Kruskal–Wallis test. Beta diversity was calculated using weighted UniFrac distances and visualized by principal coordinates analysis (PCoA). Differences in community composition were evaluated by permutational multivariate analysis of variance (PERMANOVA; 999 permutations). Differentially abundant taxa were identified using LEfSe (Linear Discriminant Analysis Effect Size) or Kruskal–Wallis testing with Benjamini–Hochberg correction, as indicated in the figure legends. Analyses were performed using QIIME and Python 3.10 (pandas, scipy, scikit-bio, statsmodels)

### scRNA-seq sample preparation

For scRNA-seq experiments, B16/F10-bearing mice were treated as described above for the corresponding *in vivo* studies. For the DCA experiment, mice received intratumoral injections of diABZI and/or DCA on days 8 and 11 after tumor inoculation, and tumors were collected on day 13. For the bacterial experiment, mice were treated with antibiotics and orally gavaged with bacteria as described above, followed by a single intratumoral injection of diABZI on day 9 after tumor inoculation; tumors were collected on day 12. Tumors were processed as described above for flow cytometry analysis. Briefly, tumors were enzymatically digested, passed through a 70 μm filter, treated with ACK red blood cell lysis buffer, and washed in cold PBS containing 2% FBS. Single-cell suspensions were stained with anti-CD45 antibody (APC-Fire 750, clone 30-F11, BioLegend, 103153), TotalSeq-C anti-mouse hashtag antibodies (BioLegend), and SYTOX Blue viability dye. Live CD45⁺ cells were sorted on a Bigfoot Spectral Cell Sorter (Thermo Fisher Scientific; 9-laser configuration) using a 100 μm nozzle at 25 psi. For each sample, 10,000 live CD45⁺ cells were sorted. Hashtagged cells from individual mice were then pooled, and 50,000 total cells were targeted for droplet encapsulation using the Chromium Single Cell platform (10x Genomics). Single-cell gene expression and feature barcode libraries were prepared according to the manufacturer’s protocol using the Chromium GEM-X Single Cell 5′ Reagent Kits v3 and Chromium GEM-X Single Cell 5′ Feature Barcode Kit v3 (10x Genomics, CG000734).

### scRNAseq analysis

Libraries were sequenced to a depth of 800M reads (GEX) and 200M reads (CSP) on an Element Aviti sequencer by the Genomics Core at Scripps Research. Reads were processed and aligned using cell ranger count (v9.0.1). Data were then analyzed using R with the “Seurat” package as well as custom scripts(*86–90*). HTO identities (that is, cells originating from each of the original 16 mice) were assigned using the *HTODemux* function, and cells without an unambiguous HTO identity or those determined to be a doublet were excluded(*91*). Then, cells with mitochondrial genes accounting for >5% or >0.4% of gene counts were eliminated. Then, the top 2,000 variable genes were identified and scaled. A principal component analysis (PCA) was performed on these genes and the top 30 principal components were used to assign k-nearest neighbors, generate a shared nearest-neighbor graph and then optimize the modularity function to determine clusters, at resolution = 0.3 (*92*). Cell type identities and clusters were then reconciled manually using prior knowledge, literature searches, and marker expression. This involved re-clustering “myeloid” and “lymphoid” clusters separately to refine marker gene expression for cell identification. Clusters were then re-named with appropriate cell type identification. Cells belonging to clusters that could not be readily identified, constituting ∼5% of cells, were eliminated from subsequent analysis. Differential gene expression analysis was carried out using default parameters of the *FindMarkers* function. The diABZI response was defined as genes induced (log2FC > 0.5 and adjusted p-value 0.05) in each cell type between DMSO and diABZI groups, and then mapped onto all cells of that cell type using the *AddModuleScore* function. All other significant gene expression changes (e.g. per cell *Csc* vs *Csp* comparison) were defined as absolute value of log_2_FC > 0.25 and adjusted p-value 0.05. KS tests were used to determine whether the diABZI response was significantly altered for each cell type across the relevant comparisons. A chi-square test was used to determine whether different cell types were over- or under-represented in the various groups in the relevant pairwise comparisons. All R scripts are available upon request.

### Statistics

Statistical analyses were conducted using GraphPad Prism 8.0. Briefly, pairwise comparisons were performed by using unpaired two-sided *t*-tests or two-sided Mann–Whitney tests. For comparisons of multiple groups with one variable, one-way analysis of variance (ANOVA) followed by multiple comparisons testing was used. For comparisons of multiple groups with two variables, two-way ANOVA followed by multiple comparisons testing was used. Adjusted *P* values are provided for multiple comparisons. Specific statistical tests are identified in the corresponding figure legends. Comparisons with *P* > 0.05 were not considered significant.

## Data Availability

The data supporting the findings of this study are available within the paper and its Supplementary Information. The raw proteomics data have been deposited in the ProteomeXchange Consortium via the PRIDE repository under accession **PXD056524**. The atomic model and cryo-EM density map generated in this study have been deposited in the Protein Data Bank and Electron Microscopy Data Bank under accession codes **9N7F** and **EMD-49094**, respectively.

## Acknowledgments

We thank the Scripps Research Flow Cytometry, Imaging, and Genomics Cores for technical support.

## Author contributions

X.Y., X.Z. and H.C.H. conceived the study. X.Y., X.Z., W.L., A.G., A.H., A.T., C.W., Y.-Z.H., A.S., S.D. and X.H.Z. designed and performed methodology, investigation and visualization. H.C.H., C.G.P., A.B.W. and L.L.L. acquired funding. H.C.H. administered the project. Supervision was provided by H.C.H., C.G.P., A.B.W., L.L.L. and S.F. X.Y., X.Z. and H.C.H. wrote the original draft, and all authors contributed to review and editing of the manuscript.

## Additional information

Supplementary Information is available for this paper.

Correspondence and requests for materials should be addressed to Howard C. Hang.

---

**Extended Data Figure 1.**
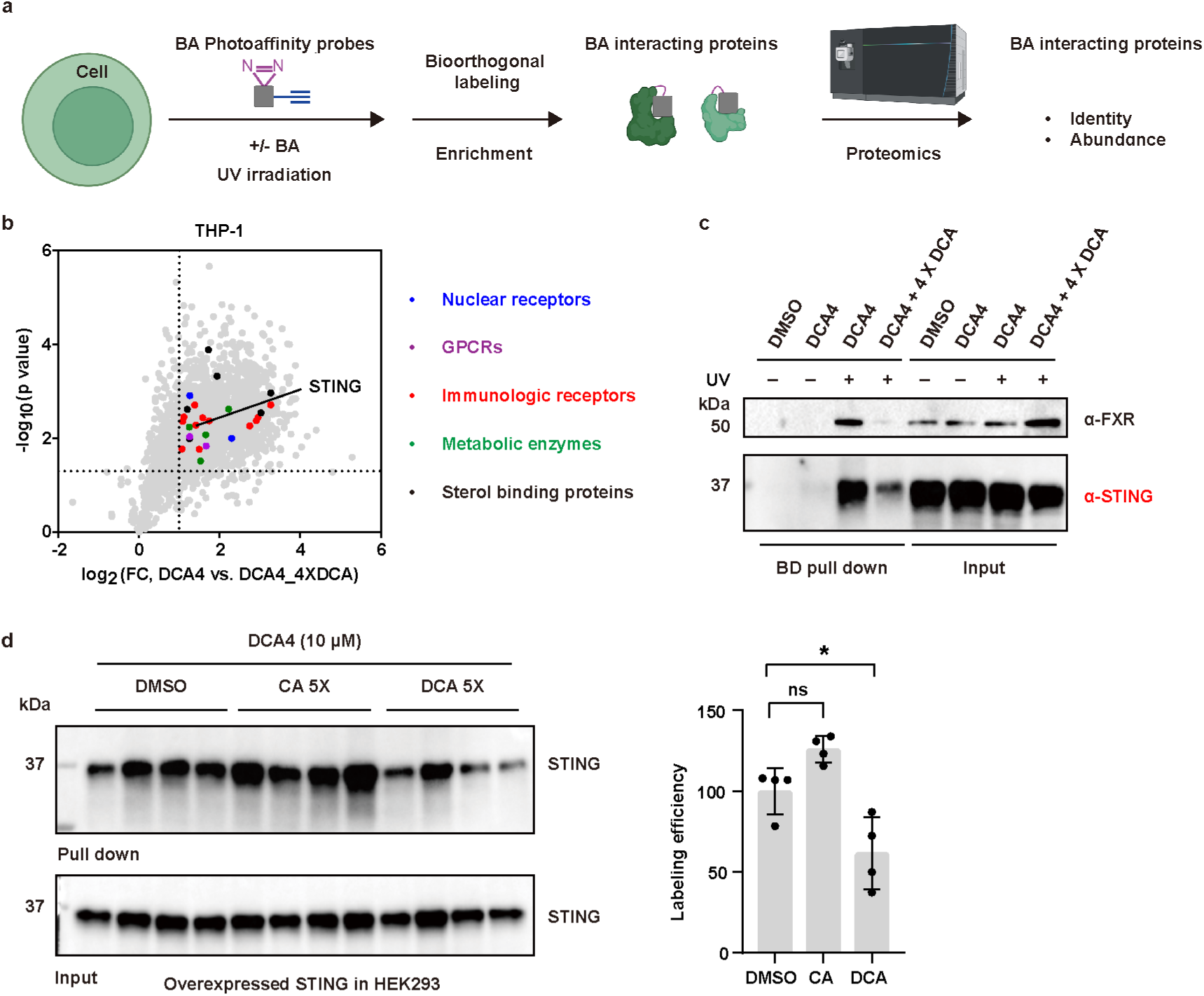
Chemoproteomic profiling of DCA-interacting proteins. (a) Schematic workflow for chemoproteomic profiling of bile acid-interacting proteins using photoaffinity probes. Cells are incubated with bile acid photoaffinity probes containing a photoreactive diazirine group and an alkyne handle. Upon UV irradiation, the diazirine generates a reactive carbene intermediate that covalently crosslinks the probe to nearby interacting proteins in live cells. Probe-labeled proteins are subsequently conjugated to biotin through copper-catalyzed azide–alkyne cycloaddition, enriched with streptavidin beads, digested into peptides, and identified by quantitative LC-MS/MS-based proteomics. Competition with excess parent bile acid is used to help distinguish specific bile acid-interacting proteins from nonspecific probe-enriched proteins. (b) DCA4-enriched proteins are competed off by excess of DCA in THP-1 cells. STING is one of proteins that are selectively enriched by DCA photoaffinity reporters. Other DCA-enriched proteins are highlighted by different colors. (c) Validation of DCA-FXR/STING interaction in HT-29 by reverse western blot. HT-29 cells were treated with bile acid photoaffinity reporters with or without excess of DCA for 0.5 h, followed by 365-nm light irradiation. The labeled proteins were reacted with az-biotin and enriched with streptavidin beads (‘pulldown’). Equal loading was validated by immunoblotting of input samples before enrichment (‘input’) (d) Validation of DCA-STING interaction in HEK293T by reverse western blot. HEK293T transfected with pMSCV-hygro-STING were treated with DCA4 in the presence or absence of 5X CA and 5X DCA for 0.5 h, followed by 365-nm light irradiation. The labeled proteins were reacted with az-biotin and enriched with streptavidin beads (‘pulldown’). Equal loading was validated by immunoblotting of input samples before enrichment (‘input’) (*n* = 4 biologically independent samples). A one-way ANOVA followed by Tukey’s multiple comparisons test was used to calculate the adjusted P values. *P < 0.05, **P < 0.01, ***P < 0.001, ****P < 0.0001; ns, not significant.

**Extended Data Figure 2.**
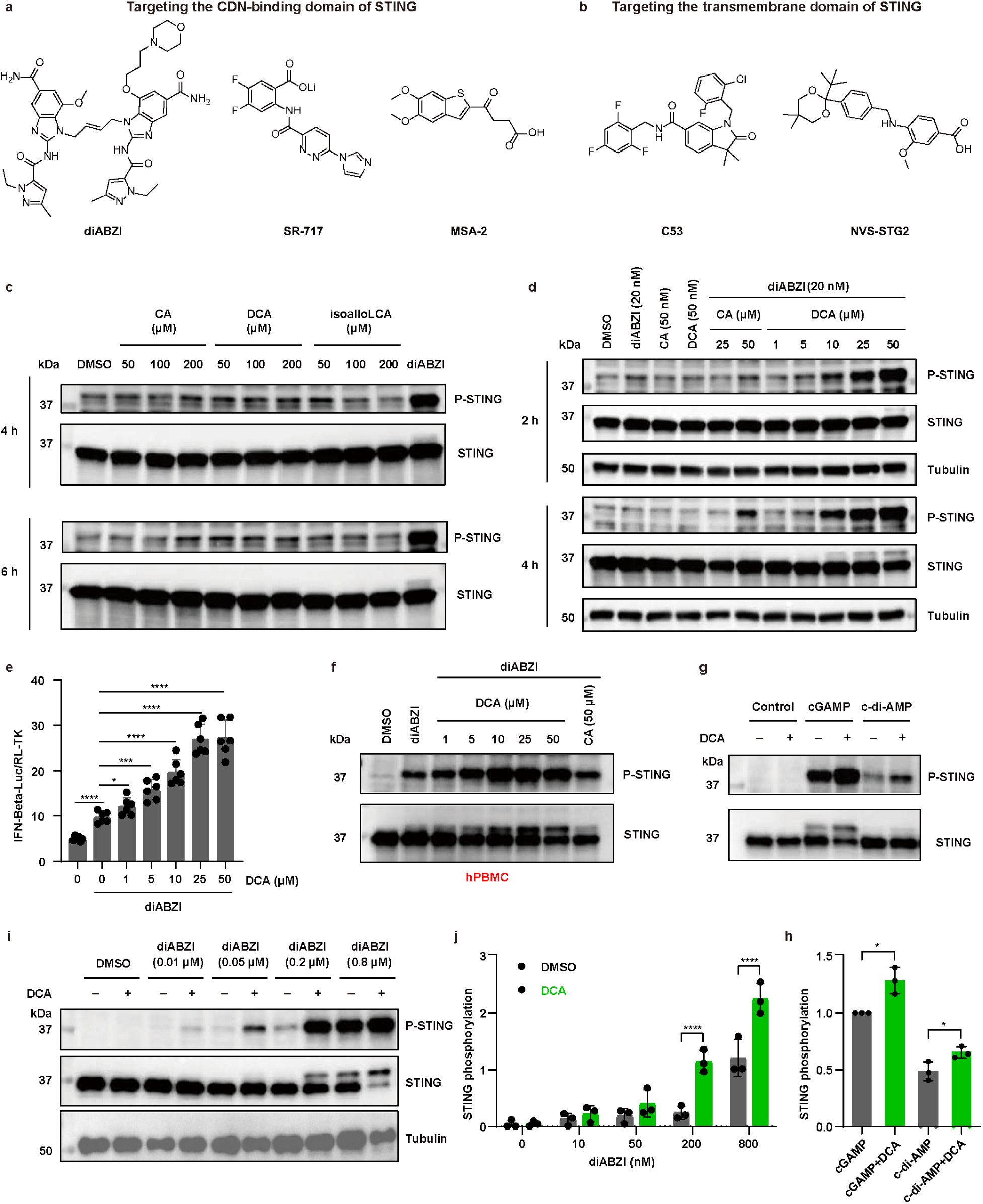
Bile acids synergize with STING agonists to promote STING activation ex vivo. (a) Chemical structure of STING agonists targeting CDNs binding domain. (b) Chemical structure of STING agonists targeting STING transmembrane domain. (c) Western blot analysis of cell lysates from THP-1 cells treated with diABZI (100 nM) and different concentrations of bile acids (50, 100, 200 μM) alone for 4 or 6 hours. (d) Western blot analysis of cell lysates from THP-1 cells treated with diABZI (20 nM), CA (25, 50 µM) and DCA (1, 5, 10, 25, 50 µM) alone or together for 2 or 4 hours. (e) HEK293T transfected with pMSCV-hygro-STING, IFN-Beta_pGL3 and pRL-TK control plasmid were treated with diABZI (20 nM) in the presence or absence of different doses of DCA (*n* = 6 biologically independent samples). Data were analyzed by unpaired two-sided t-test. The apparent EC₅₀ of DCA for enhancing diABZI-induced IFN-β reporter activation was 9.36 μM. (Dose–response curves were fitted using nonlinear regression in GraphPad Prism with a four-parameter variable-slope logistic model. The EC_50_ was calculated from the best-fit curve.) (f) Western blot analysis of cell lysates from Human PBMCs treated with diABZI (200 nM) in the presence or absence of DCA and CA for 2 hours. (g) Western blot analysis of cell lysates from Human PBMCs treated with STING agonists (cGAMP (5 µg/mL), c-di-AMP (5 µg/mL), transfected with lipo2000 (1 µg/µL)) and DCA (50 µM) alone or together for 1 hour. (i) Western blot analysis of cell lysates from Human PBMCs treated with different concentrations of diABZI with or without DCA (50 µM) for 2 hours. (h, j) Quantification of phosphorylated STING by grayscale analysis (p-STING/STING) (n = 3 biologically independent samples). Tubulin is shown as an additional loading control. For e, data were analyzed by unpaired two-sided t-test. for g, a two-way ANOVA followed by Sidak’s multiple comparisons test was used to calculate adjusted P values. *P < 0.05, ****P < 0.0001.

**Extended Data Figure 3.**
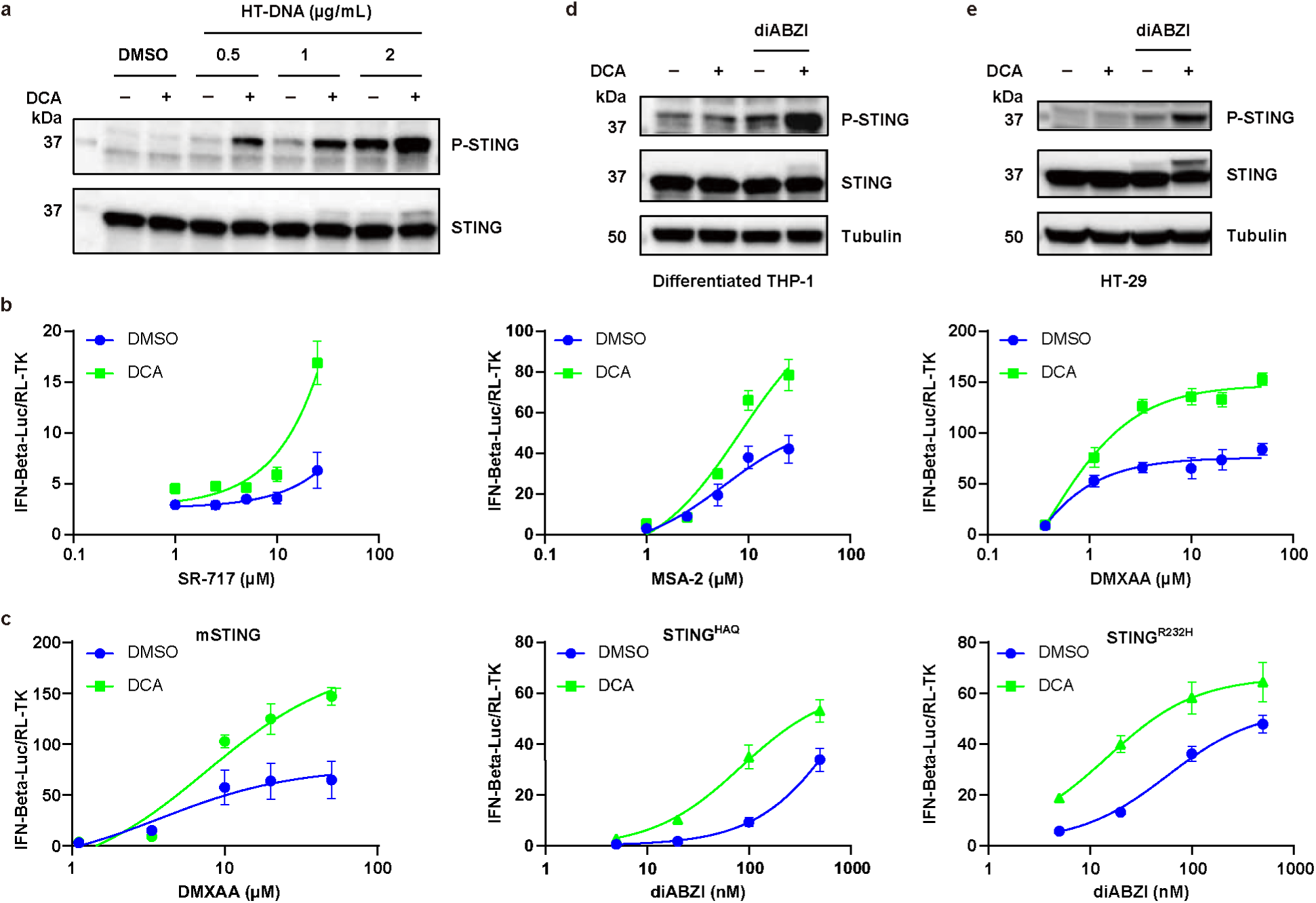
The generality of DCA activity on STING activation. (a) Western blot analysis of cell lysates from differentiated THP-1 cells (Cells were pretreated with 100 nM of PMA for 24 h) treated with different concentrations of HT-DNA (0.5, 1, 2 μg/mL, HT-DNA was transfected by Lipo2000 (1 µg/µL)) in the presence or absence of DCA (50 µM). (b) HEK293T cells transfected with pMSCV-hygro-STING, IFN-Beta_pGL3 (a luciferase reporter plasmid under control of the IFN-β promoter) and pRL-TK control plasmid were treated with different STING agonists (SR-717, MSA-2 and DMXAA) in the presence or absence of DCA (50 µM). For DMXAA, pMSCV-hygro-STING(S162A, G230I, Q266I) was used (*n* = 3 biologically independent samples). (c) HEK293T transfected with different STING variants (pMSCV-hygro-murineSTING, pMSCV-hygro-STING HAQ and pMSCV-hygro-STING R232H), IFN-Beta_pGL3 (a luciferase reporter plasmid under control of the IFN-β promoter) and pRL-TK control plasmid were treated with STING agonist (DMAXX or diABZI) in the presence or absence of DCA (50 µM) (For mSTING, *n* = 3, for STING^HAQ^, *n* = 5, for STING ^H232^, *n* = 4, biologically independent samples). (d) Western blot analysis of cell lysates from differentiated THP-1 cells (pretreat the cells with 100 nM of PMA for 24 h) treated with diABZI (50 nM) in the presence or absence of DCA (50 µM). (e) Western blot analysis of cell lysates from HT-29 cells treated with diABZI (50 nM) in the presence or absence of DCA (50 µM).

**Extended Data Figure 4.**
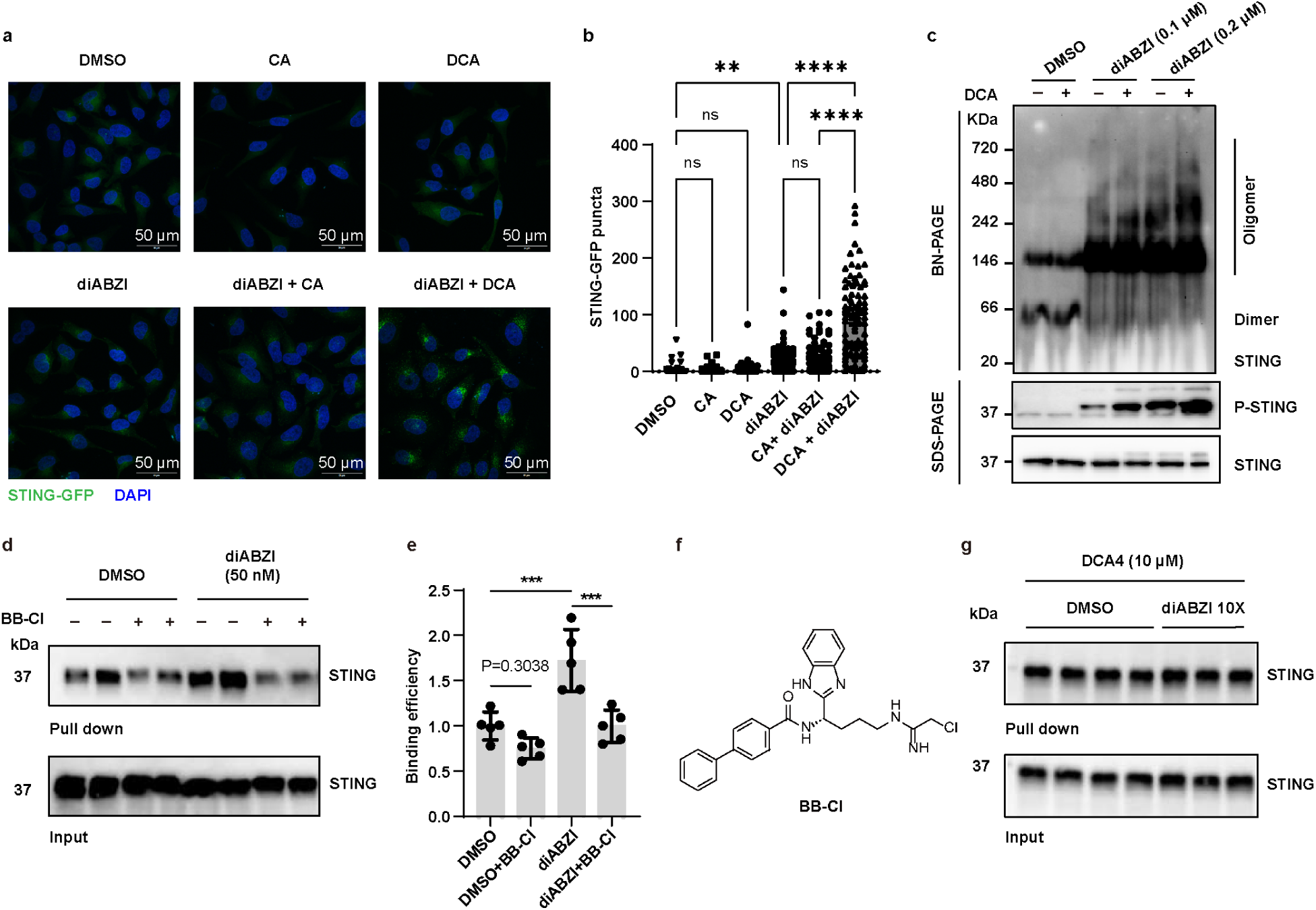
DCA promotes diABZI-induced STING oligomerization. (a) Representative images of HeLa cells stably expressing STING-GFP. Cells were treated with CA (50 µM) or DCA (50 µM), in the presence or absence of diABZI (50 nM), using equal volume of DMSO as a control. Localization of STING-GFP in cells was monitored by the fluorescence signal of GFP puncta. Nuclei were counter-stained with DAPI. The experiments were repeated three times. The images are representative images from these experiments. (b) Quantification of intracellular STING puncta. The bar graph shows the amount of STING puncta per cell from each treatment group. Data represent mean ± SD and were analyzed by the one-way ANOVA with Šídák’s multiple comparisons test. *P < 0.05, **P < 0.01, ***P < 0.001, ****P < 0.0001; ns, not significant. Scale bar, 50 µm. (c) STING oligomerization induced by DCA, diABZI or both. THP1-Dual cells were stimulated with DCA (50 μM) in the presence or absence of diABZI (100, 200 nM) for 2 hours. STING oligomerization was analyzed by Blue native PAGE (BN-PAGE), and the indicated proteins were detected by immunoblotting. (d) THP-1 cells were pretreated with diABZI (50 nM) in the presence or absence of BB-Cl (10 µM) for 1 hour. Cells were then treated with DCA4 (25 µM) for another 0.5 hour, followed by 365-nm light irradiation. The labeled proteins in cell lysis were reacted with az-biotin and enriched with streptavidin beads (‘pulldown’). Equal loading was validated by immunoblotting of input samples before enrichment (‘input’). (e) Quantification of DCA4 STING binding efficiency in d by grayscale analysis (pull down/input) (*n* = 3 biologically independent samples). A one-way ANOVA followed by Tukey’s multiple comparisons test was used to calculate the adjusted P values. (f) Chemical structure of an inhibitor of STING activation, BB-Cl. (g) Validation of DCA-STING interaction in HEK293T overexpressed STING in the presence or absence of excess of diABZI by reverse western blot. HEK293T transfected with pMSCV-hygro-STING were treated with DCA4 in the presence or absence 10X diABZI for 0.5 h at 37 °C, followed by 365-nm light irradiation. The labeled proteins were reacted with az-biotin and enriched with streptavidin beads (‘pulldown’). Equal loading was validated by immunoblotting of input samples before enrichment (‘input’). *P < 0.05, **P < 0.01, ***P < 0.001, ****P < 0.0001; ns, not significant.

**Extended Data Figure 5.**
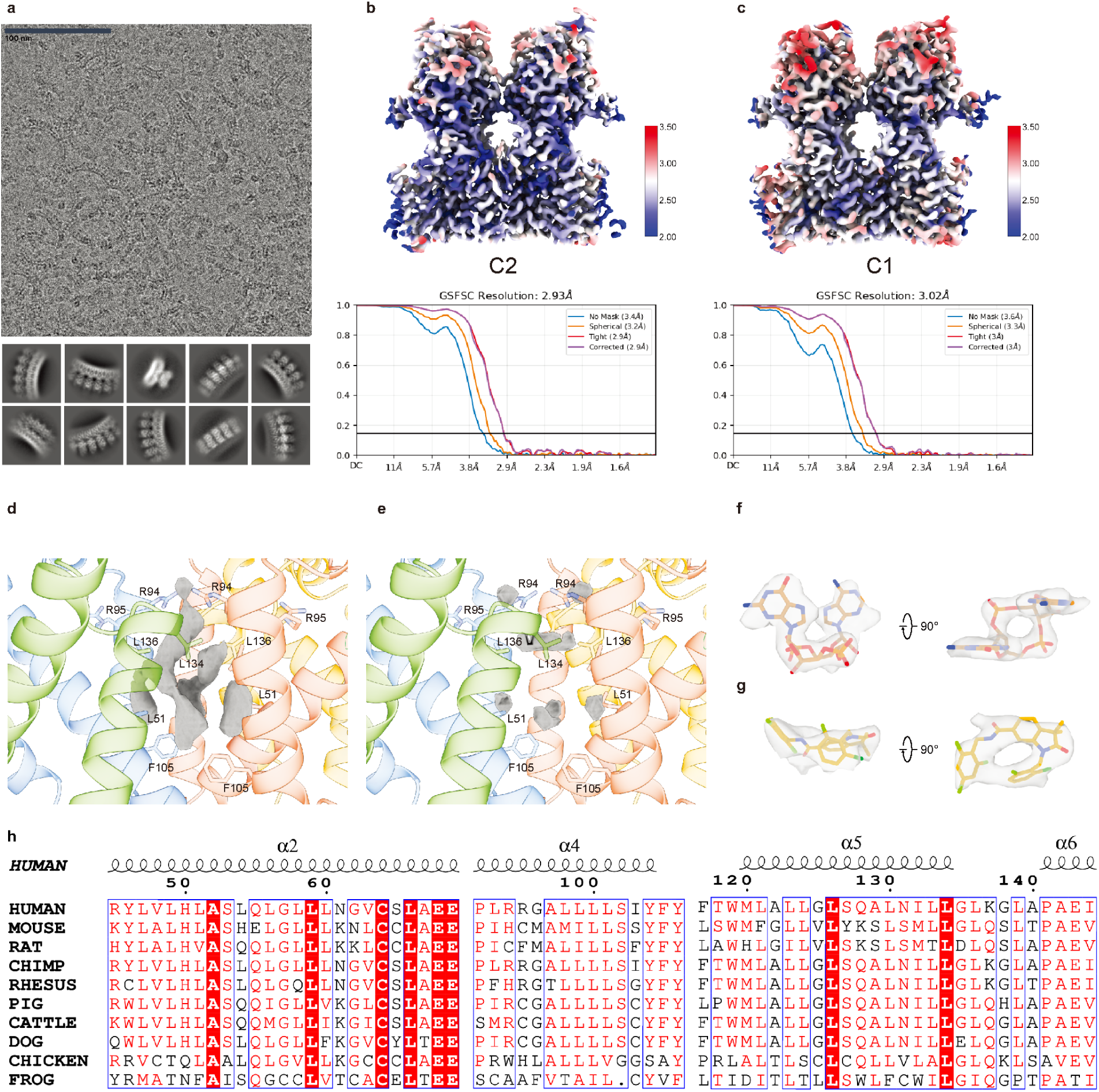
Cryo-EM analysis of STING binding with different agonists. (a) Motion corrected micrograph and 2D class averages of high-order oligomers of human STING. (b,c) 3D reconstruction of the STING tetramer complexed with ligands (cGAMP and C53), and gold-standard FSC curve of the final 3D reconstruction. (d,e) High-resolution cryo-EM map of the two STING dimer with highlight of dimer-dimer interface molecular density. (d: STING, cGAMP, C53 and DCA (C1 symmetry, this study); e: STING, cGAMP, C53 (PDB ID: 7SII)) (f) Cryo-EM density for cGAMP. (g) Cryo-EM density for C53. (h) Evolutionary conservation analysis of the putative DCA-associated STING transmembrane region. Multiple sequence alignment of representative vertebrate STING orthologs around the proposed DCA-associated transmembrane region. Secondary-structure elements are indicated above the alignment and residues are numbered according to human STING. Conserved residues are shown in white letters on a red background, and similar residues are shown in red letters. The alignment was generated using ClustalW and rendered with ESPript.

**Extended Data Figure 6.**
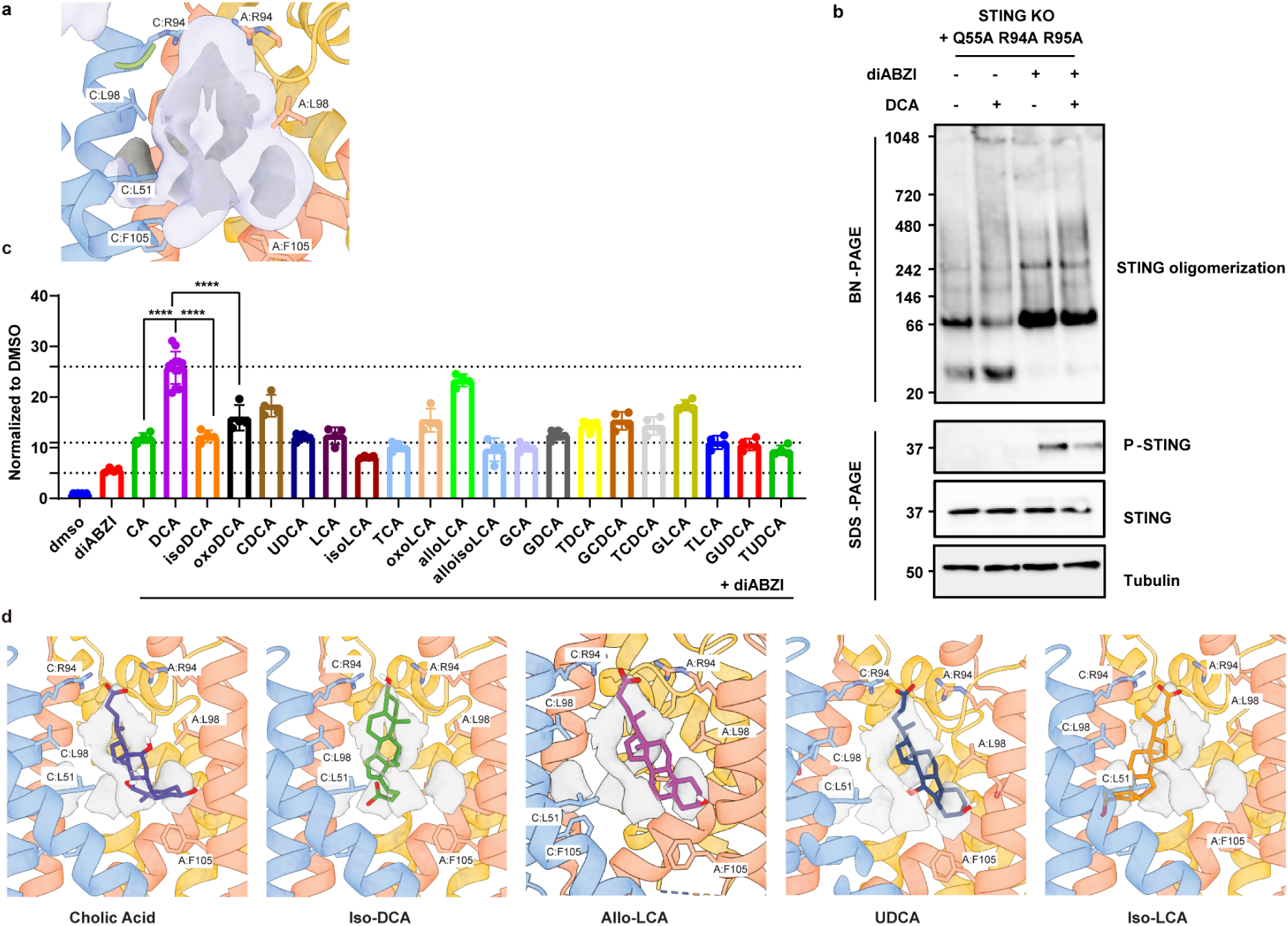
Responses of human STING to different bile acids, with key binding pocket mutations. (a) Illustration of docking density cloud (outer violet surface), representing Boltzmann-weighted map of ligand states sampled during docking simulation. All experimental density lobes on one side of the binding pocket were sampled during the simulation, but less so on the opposite side of the pocket. (b) STING oligomerization induced by DCA, diABZI or both. THP1-Dual STING KO cells (STING Q55A+R94A+R95A stable expression cell line) were stimulated with DCA in the presence or absence of diABZI for 2 hours. STING oligomerization was analyzed by Blue native PAGE (BN-PAGE), and the indicated proteins were detected by immunoblotting. (c) HEK293T transfected with pMSCV-hygro-STING, IFN-Beta_pGL3 (a luciferase reporter plasmid under control of the IFN-β promoter) and pRL-TK control plasmid were treated with diABZI (20 nM) in the presence or absence of different conjugated and free bile acids (50 µM) (For DCA, *n* = 12, and others *n* = 4, biologically independent samples). Data were analyzed by unpaired two-sided t-test. (d) Docked models of Cholic Acid, Iso-DCA, UDCA, Iso-LCA and alloLCA in STING tetramer. Dockings guided by C2-symmetric density for DCA, shown in gray. Cholic acid, UDCA and alloLCA adopt binding poses like that predicted for DCA. However, alloLCA lacks an additional hydroxyl group that may stabilize the STING tetramer more like DCA. Iso-DCA, meanwhile, is the only ligand to orient the carboxylate down into the bottom of the pocket, with a hydroxyl hydrogen bonding with R94 of the C monomer. Iso-LCA also adopts a different binding mode from cholic acid. While Iso-LCA does place the carboxylate in a salt bridge with R94 of the A and C monomers, the rest of the molecule is oriented with interactions with the C monomer. All static poses for all of the tested bile acids were consistent across simulation runs. *P < 0.05, **P < 0.01, ***P < 0.001, ****P < 0.0001.

**Extended Data Figure 7.**
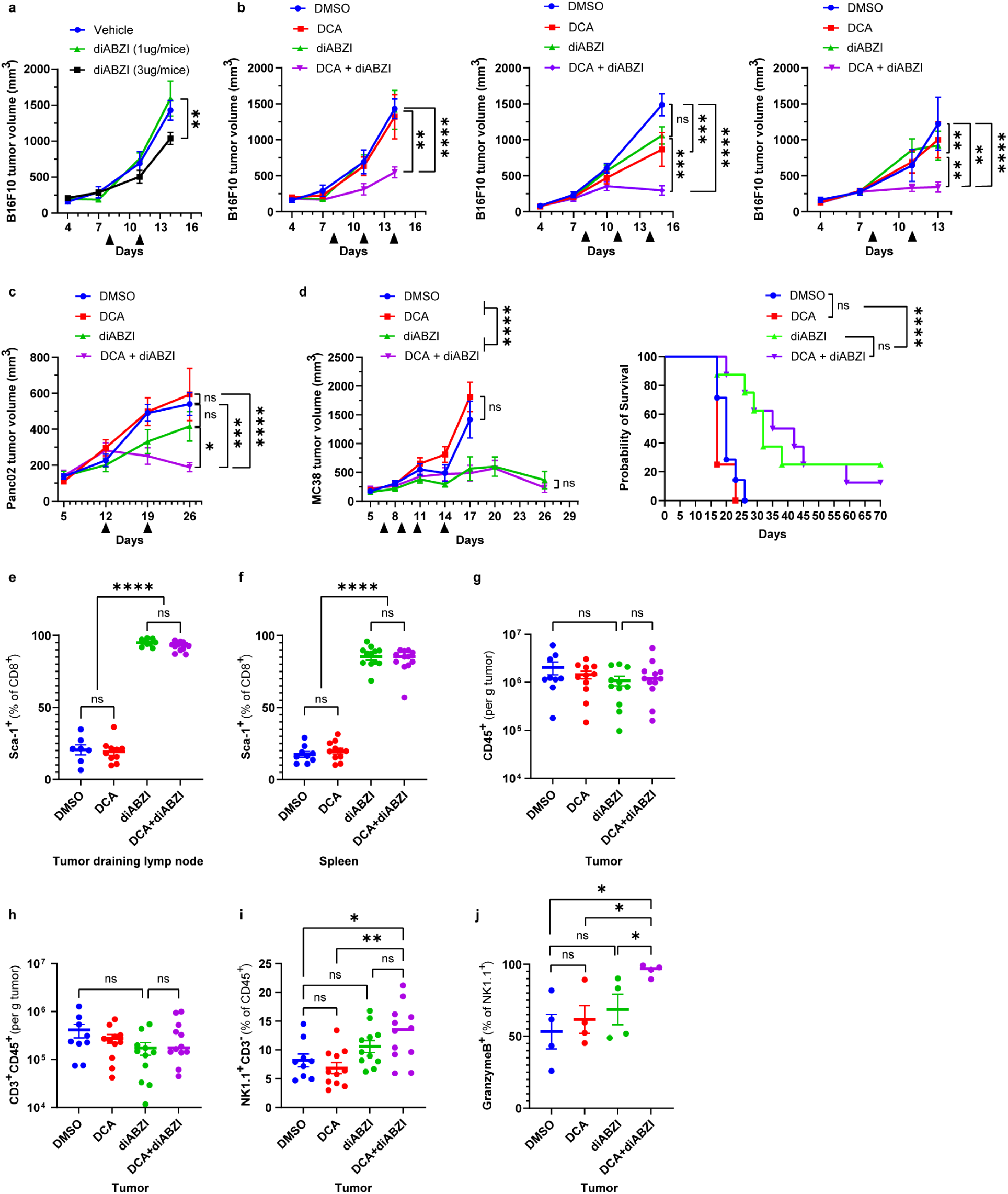
The effect of DCA on STING-mediated antitumor activity in different tumor models. (a) Comparison of B16-F10 tumor growth following i.t. injection of two different concentrations of diABZI. (b) All the other repeats for B16-F10 melanoma tumor experiments. (c) Panc02 tumor growth in mice treated i.t. with vehicle (DMSO), DCA, diABZI, or the combination. (d) MC38 tumor growth and survival in SPF mice that were IT injected with vehicle (DMSO), bile acids DCA, diABZI or the combination (DCA+ diABZI). Arrowhead indicates the day for i.t. injection. *n* = 8 mice per group. Data represent mean ± SEM and were analyzed using a mixed effects model with Tukey’s multiple comparisons post hoc test. **P < 0.01, ***P < 0.001, ****P < 0.0001; ns, not significant. (e, f) Percentage of Sca-1^+^ cells among CD8⁺ T cells in the tdLN (e) and spleen (f). (g) Quantification of tumor infiltrating CD45^+^ cells. (h) Quantification of tumor infiltrating CD3^+^ CD45^+^ cells. (i) Quantification of tumor infiltrating NK1.1^+^ CD3^-^ cells. (j) Quantification of tumor infiltrating Granzyme B^+^ NK1.1^+^ cells. Data for a-e were pooled from two independent experiments. *n* = 9-12 per group. Data for f represents a single representative experiment, with *n* = 4 per group. All the data represent mean ± SEM and were analyzed by the unpaired t test. *P < 0.05, **P < 0.01, ****P < 0.0001; ns, not significant.

**Extended Data Figure 8.**
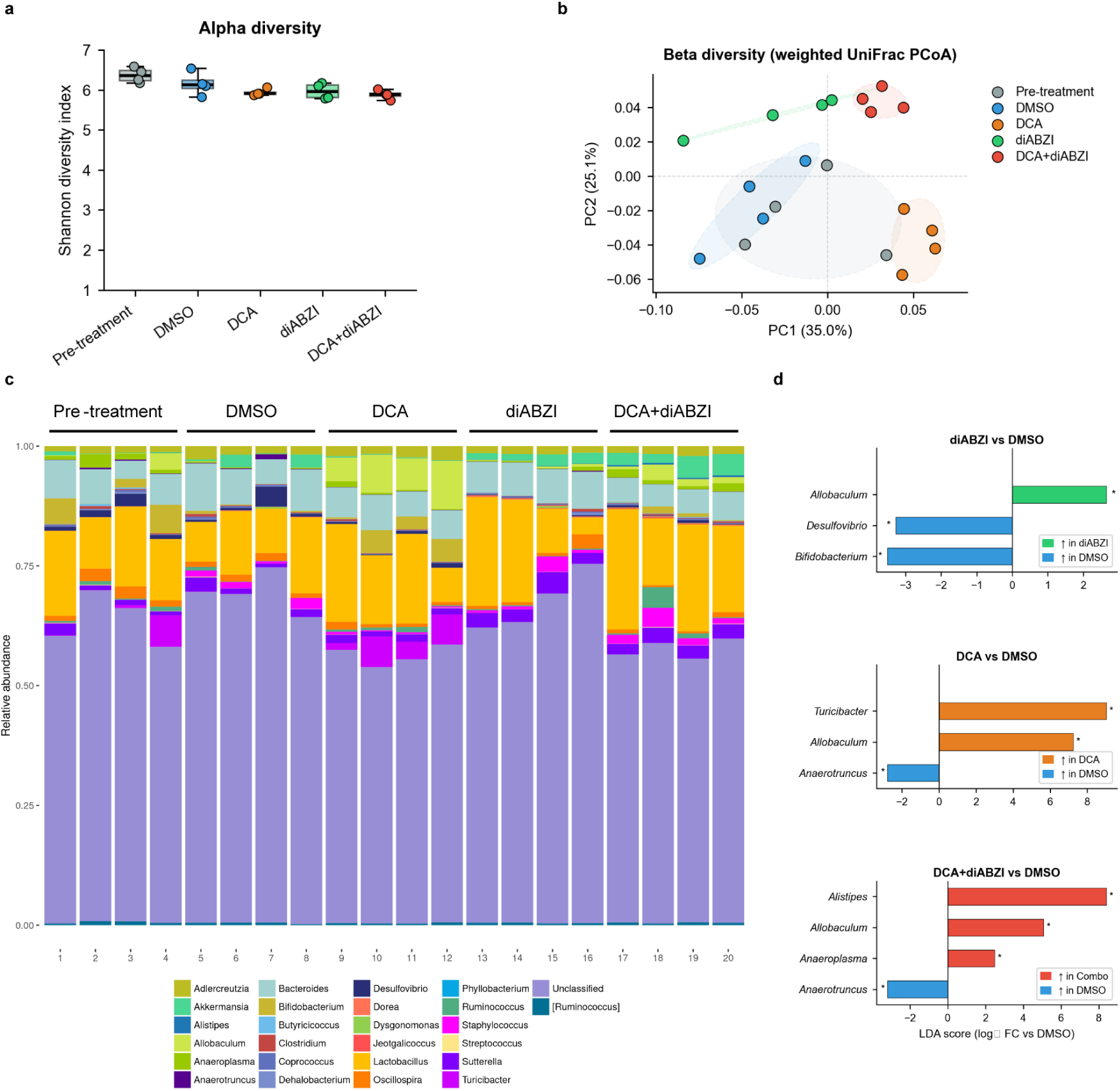
16S rRNA microbiome profiling reveals microbiota shifts without enrichment of innate immune-activating taxa. (a) Alpha diversity (Shannon index) across treatment groups. n = 4 mice per group. Statistical comparison by Kruskal-Wallis test. (b) Beta diversity by weighted UniFrac PCoA. Each point represents one mouse (n = 4 per group; colours as indicated). Group separation assessed by PERMANOVA (999 permutations). (c) Genus-level relative abundance. Each bar represents one mouse (n = 4 per group). (d) Differential abundance analysis (LEfSe-compatible). Log₂ fold change versus DMSO for genera with ≥0.2% mean relative abundance in at least one group. *p < 0.05, Wilcoxon rank-sum test, Benjamini–Hochberg FDR correction.

**Extended Data Figure 9.**
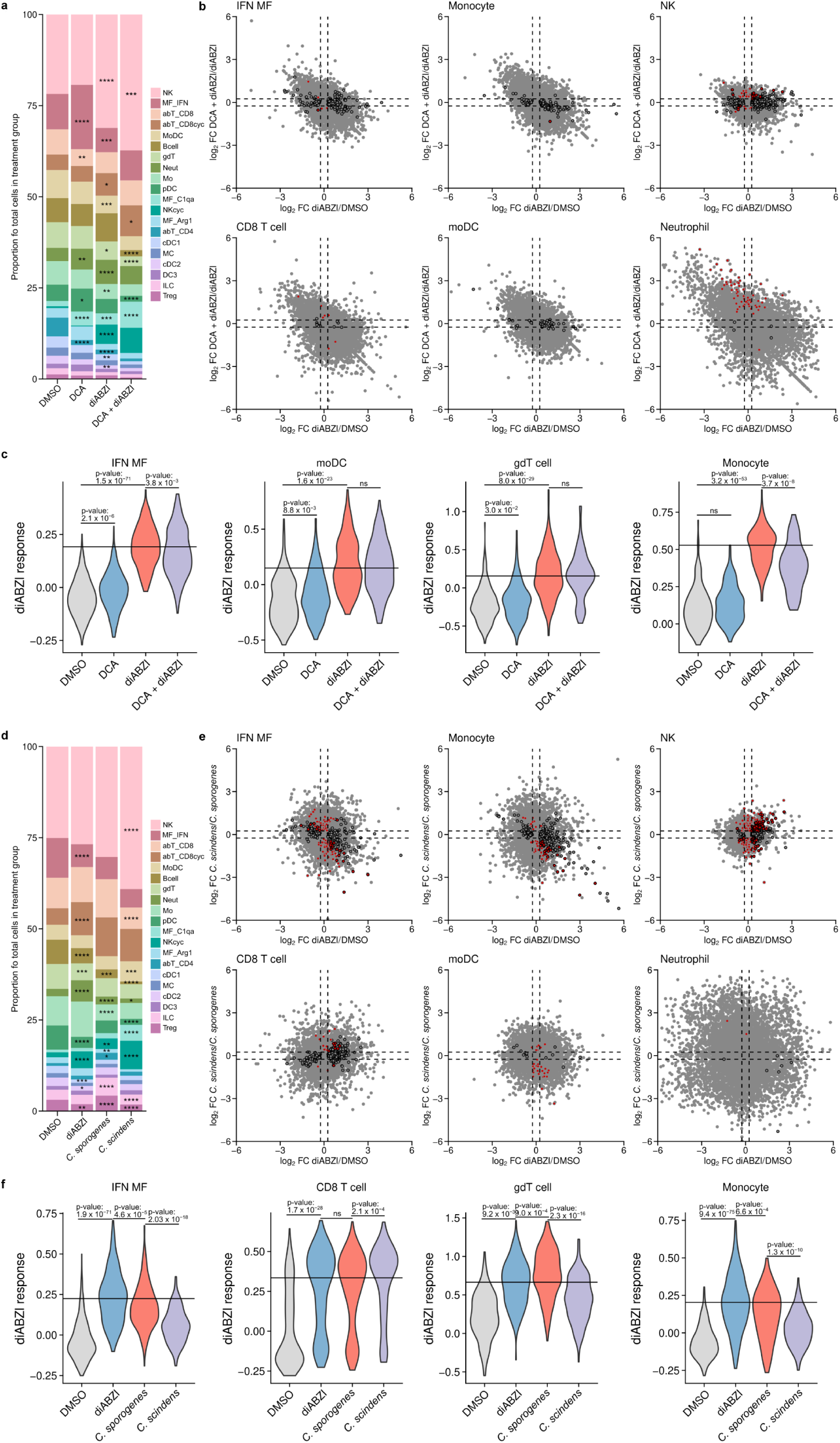
DCA and the DCA-producing bacterium *Clostridium scindens* alter tumor-infiltrating immune-cell composition and enhance diABZI-induced transcriptional responses. CD45^+^ immune cells were isolated from tumors of mice subjected to the indicated treatments (n = 4 mice per group) and analyzed by single-cell RNA sequencing (scRNA-seq). **a–c**, Effects of DCA treatment. **a**, Stacked bar plots showing the proportion of each immune-cell type within each treatment group. Asterisks indicate differences in cell-type proportions for DCA versus DMSO (DCA column), diABZI versus DMSO (diABZI column), or DCA + diABZI versus diABZI (DCA + diABZI column), determined using chi-square tests followed by Holm correction for multiple comparisons. **b**, Comparison of gene-expression changes induced by diABZI versus DMSO (x axis) and DCA + diABZI versus diABZI (y axis) within the indicated cell types. Cell types with more than ten differentially expressed genes in either comparison are shown. Genes with significant differential expression in the diABZI-versus-DMSO comparison are outlined in black, whereas genes with significant differential expression in the DCA + diABZI-versus-diABZI comparison are filled in red. **c**, Violin plots showing per-cell diABZI-response scores in the indicated cell types and treatment groups. For each cell type, the score was calculated from genes induced by diABZI treatment relative to DMSO treatment (adjusted P < 0.05 and log2 fold change > 0.5). **d–f**, Effects of colonization with *C. scindens*, with the non-DCA-producing bacterium *Clostridium sporogenes* used as a control. **d**, Stacked bar plots showing the proportion of each immune-cell type within each treatment group. Asterisks indicate differences in cell-type proportions for diABZI versus DMSO (diABZI column), *C. sporogenes* versus diABZI (*C. sporogenes* column), or *C. scindens* versus *C. sporogenes* (*C. scindens* column), determined using chi-square tests followed by Holm correction for multiple comparisons. **e**, Comparison of gene-expression changes induced by diABZI versus DMSO (x axis) and *C. scindens* versus *C. sporogenes* (y axis) within the indicated cell types. Cell types with more than ten differentially expressed genes in either comparison are shown. Genes with significant differential expression in the diABZI-versus-DMSO comparison are outlined in black, whereas genes with significant differential expression in the *C. scindens*-versus-*C. sporogenes* comparison are filled in red. **f**, Violin plots showing per-cell diABZI-response scores in the indicated cell types and treatment groups. P values in **c** and **f** were determined using Kolmogorov–Smirnov tests. For **a** and **d**, *P < 0.05; ***P < 0.001; ****P < 0.0001; ns, not significant.

**Extended Data Figure 10.**
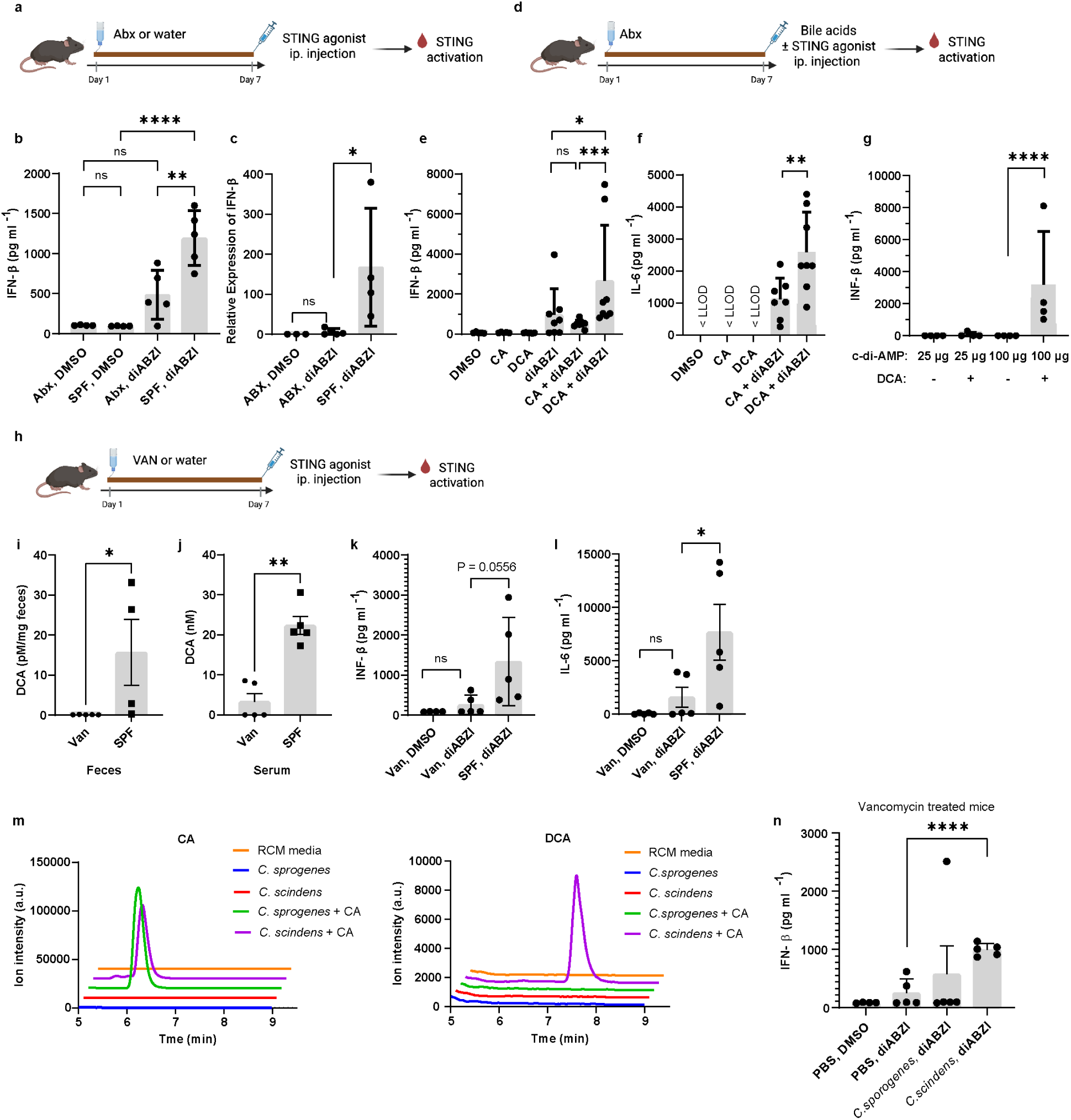
DCA and DCA-producing bacteria enhance STING-mediated type I interferon signaling *in vivo*. (a-c) Mice were treated with or without Abx for 1 week. On Day 7, Mice received an i.p. injection of diABZI. Serum IFN-β was quantified by ELISA (b), and cecal *Ifnb1* gene expression was analyzed by qPCR (c). (d-g) Mice were treated with Abx for 1 week, then i.p. injected on Day 7 with diABZI (e and f) or c-di-AMP (g) in the presence or absence of bile acids. Four hours post-injection, serum IFN-β (e and g) and IL-6 (f) levels were measured by ELISA. (h-l) Mice were treated with or without vancomycin for 1 week. On Day 7, fecal (i) and serum (j) DCA levels were measured by LC-MS. Four hours following diABZI injection, serum IFN-β (k) and IL-6 (l) were quantified by ELISA. (m) Extracted ion chromatograms (EIC) showing CA (m/z = 407.2) and DCA (m/z = 391.2) from bacterial cultures grown in RCM media with or without added CA. (n) Mice were treated with vancomycin for 7 days. On Day 7, vancomycin was replaced with water, and mice were orally gavaged daily with *C. sporogenes* or *C. scindens* for 3 consecutive days. On Day 10, mice were i.p. injected with diABZI. Four hours later, serum was collected for IFN-β quantification by ELISA. *n* = 5-8 mice per group. Data are presented as mean ± SEM and analyzed by an unpaired two-tailed *t*-test. *P < 0.05; ***P < 0.001; ****P < 0.0001; ns, not significant.

