## Supplemental file for "Microbiota-derived secondary bile acids promote STING activation and antitumor activity"

#### The PDF file includes:

Supplementary Fig. 1-2
Tables S1 to S4
Bile acid photoaffinity probe characterization

#### Other Supplementary Materials for this manuscript include the following:

Data S1
Data S2

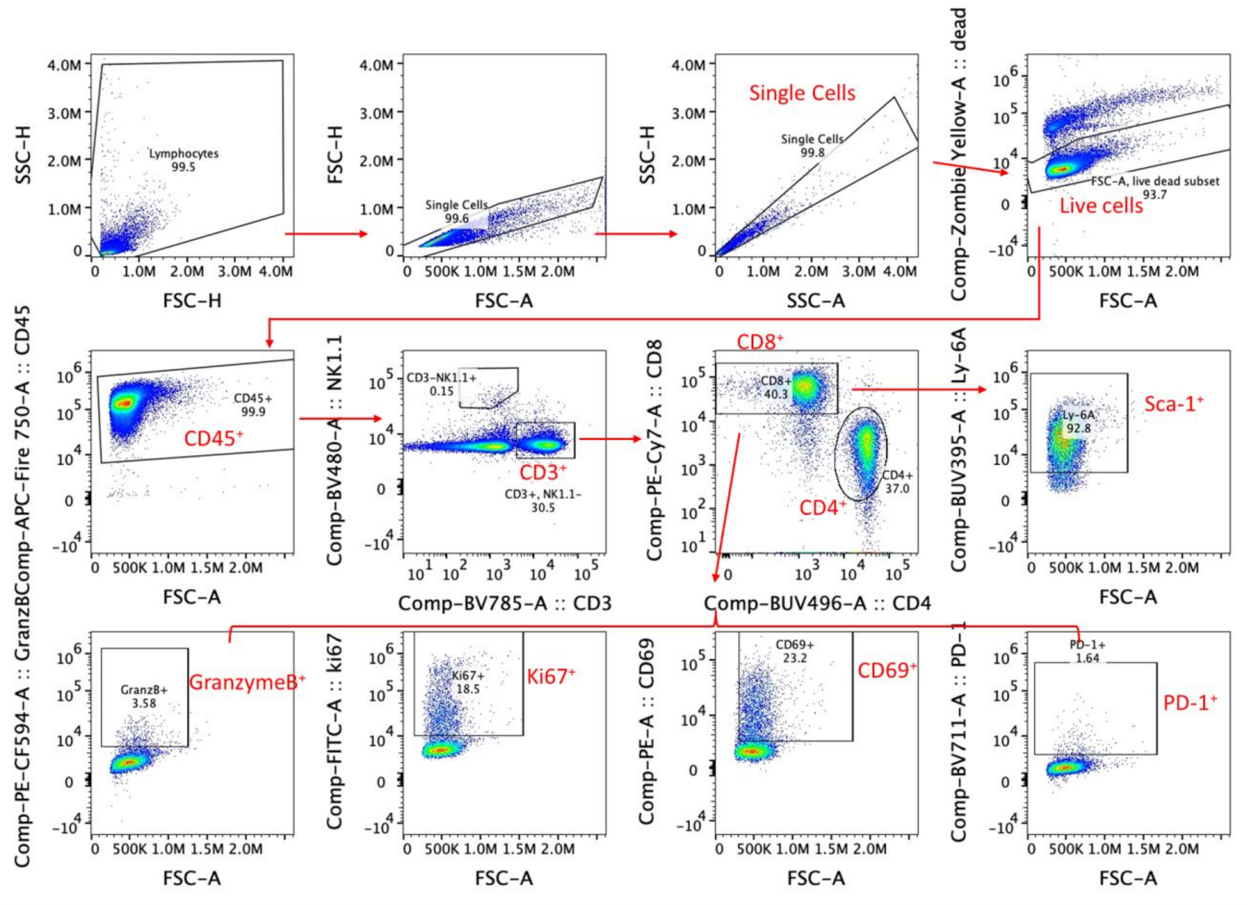

**a**. Representative plots of cells harvested from tdLN

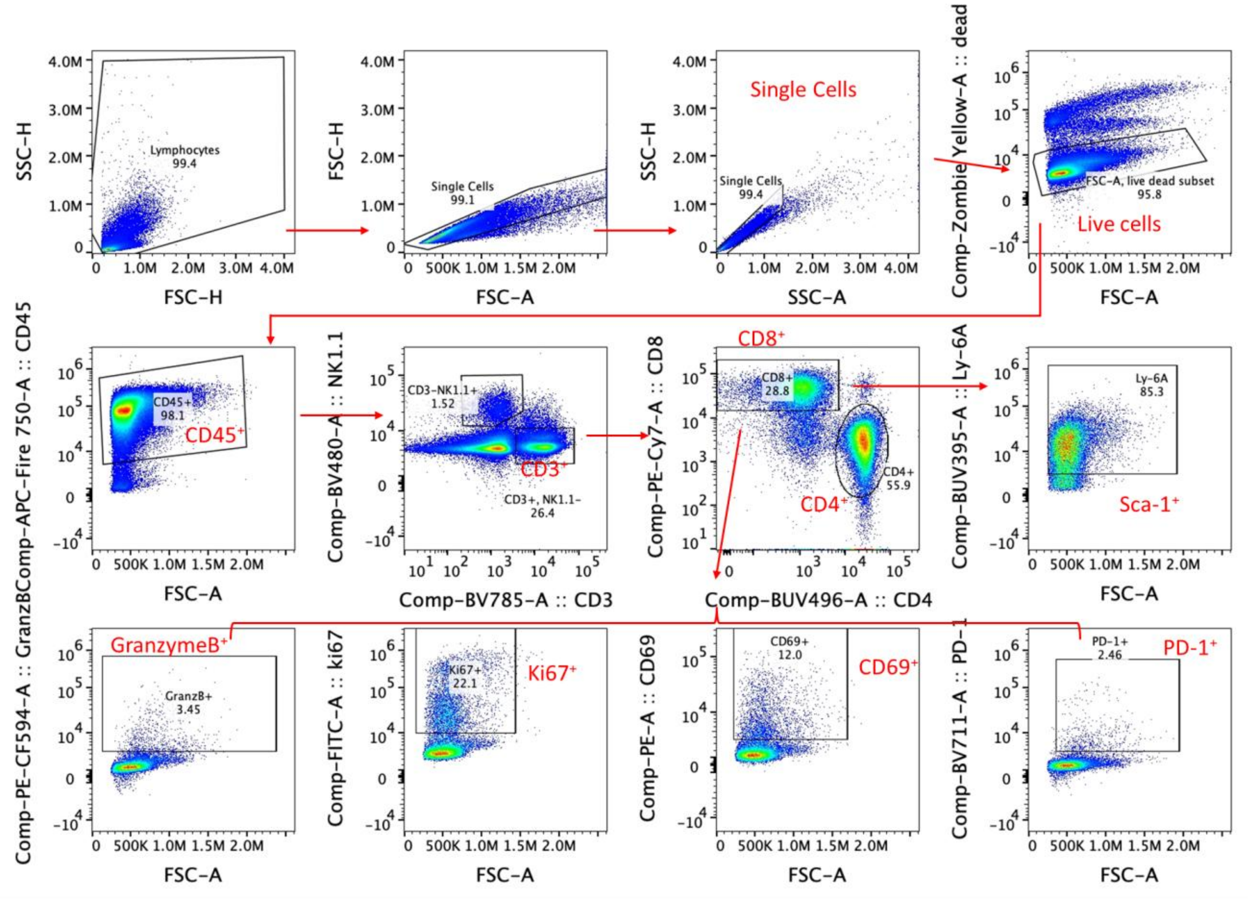

**b**. Representative plots of cells harvested from spleen

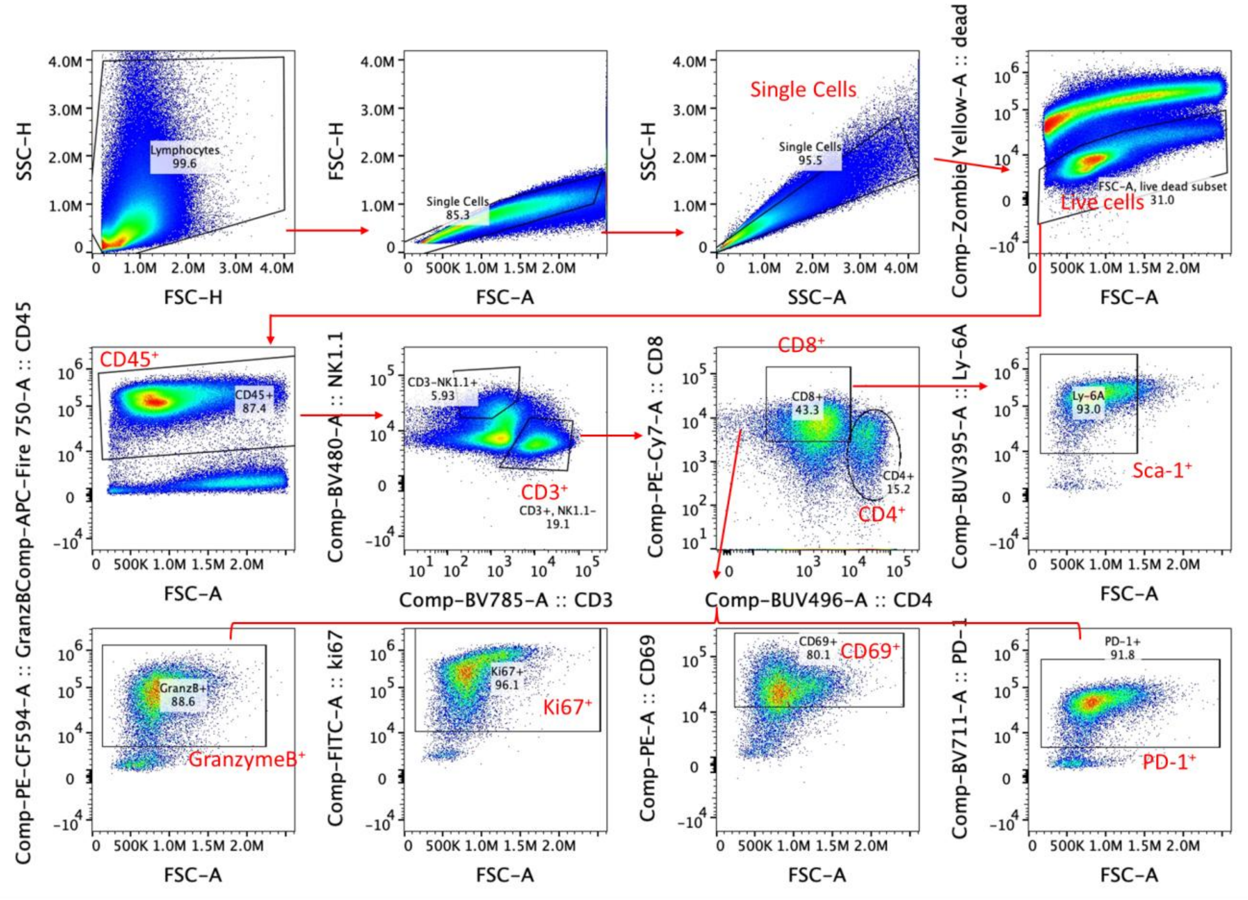

**c**. Representative plots of cells harvested from Tils

**Supplementary Fig. 1. Gating strategy used for flow cytometry analysis.** Representative plots of cells harvested from Tumor Infiltrating Lymphocytes (TILs). Single cells were selected based on forward scatter area (FSC-A) versus forward scatter height (FSC-H) parameters, as well as side scatter area (SSC-A) versus side scatter height (SSC-H) parameters. Live cells were subsequently selected from the single-cell population by gating on LIVE/DEAD dye events. Total leukocytes were identified from the live cell population using the pan-leukocyte marker CD45. T cells and NK cells were distinguished from the total leukocyte population using the pan-T cell marker CD3 and NK1.1. T cells were further subdivided into $CD4^+$ and $CD8^+$ populations based on CD4 and CD8 expression. Within the total $CD8^+$ T cell population, Sca-$1^+$ $CD8^+$ T cells, Granzyme B $^+$ $CD8^+$ T cells, Ki67 $^+$ $CD8^+$ T cells, CD69 $^+$ $CD8^+$ T cells and PD-1 $^+$ $CD8^+$ T cells were gated based on the fluorescence minus one (FMO) control.

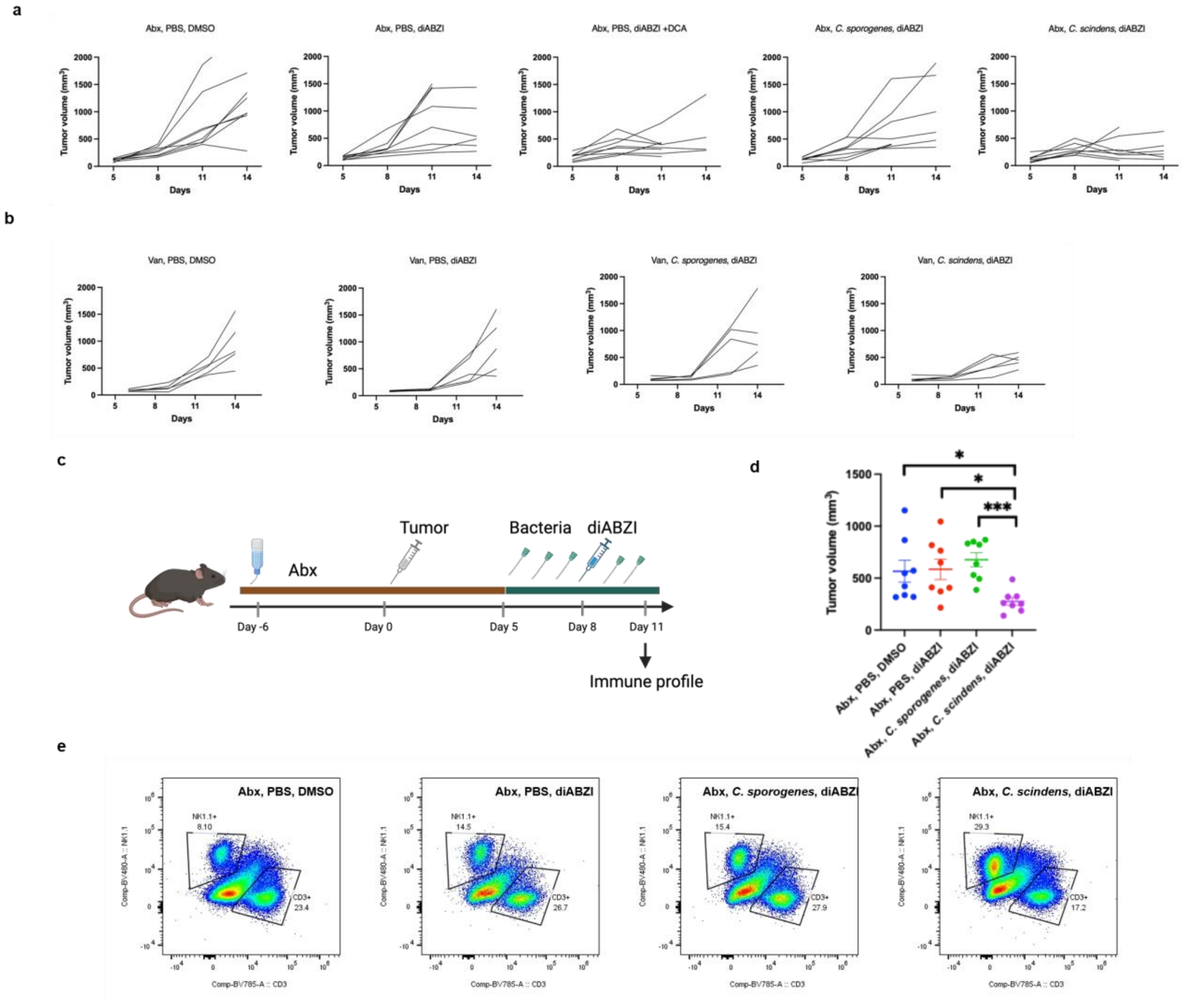

**Supplementary Fig. 2. Additional analysis of DCA-producing bacteria in STING-mediated antitumor immunity.** (a) Individual tumor growth curves for mice shown in Fig. 5b. (b) Individual tumor growth curves for mice shown in Fig. 5c. (c, d) Schematic of the tumor model (c) and tumor volumes on the day of tumor collection (d) for flow cytometric analysis shown in Fig. 5e–j. (e) Representative plots of single cell suspensions. The displayed flow-cytometry plots are representative of one individual mouse per group and are not concatenated. Complete per-mouse quantification, with each point representing one mouse, is shown in Fig. 5e–j. Graphs depict NK1.1$^+$ CD3$^-$ and NK1.1$^-$ CD3$^+$ tumor infiltrating cells.

Table S1. Highlighted proteins in Fig. 1c.

| Accession | Description | Protein | Functions | $\log_2$ Fold Change | $-\log_{10}$(P value) |
|---|---|---|---|---|---|
| Q86WV6 | Stimulator of interferon genes protein | STING | Immunology related proteins | 2.760744 | 1.985622 |
| O60603 | Toll-like receptor 2 | TLR2 | | 2.479786 | 2.188568 |
| P57764 | Gasdermin-D | GSDMD | | 1.850415 | 2.216353 |
| Q86UT6 | NLR family member X1 | NLRX1 | | 3.092457 | 2.613049 |
| P01730 | T-cell surface glycoprotein CD4 | CD4 | | 2.312721 | 2.630891 |
| Q03518 | Antigen peptide transporter 1 | TAP1 | | 3.397265 | 2.015217 |
| Q03519 | Antigen peptide transporter 2 | TAP2 | | 2.65486 | 1.749356 |
| O15533 | Tapasin | TAPBP | | 2.56642 | 2.923959 |
| Q14258 | E3 ubiquitin/ISG15 ligase TRIM25 | TRIM25 | | 1.42366 | 1.631242 |
| Q06187 | Tyrosine-protein kinase BTK | BTK | | 1.320199 | 1.380474 |
| P30273 | High affinity immunoglobulin epsilon receptor subunit gamma | FCER1G | | 2.819583 | 2.115554 |
| Q92499 | ATP-dependent RNA helicase DDX1 | DDX1 | | 1.463202 | 1.618586 |
| Q7Z2W4 | Zinc finger CCCH-type antiviral protein 1 | ZC3HAV1 | | 1.537337 | 1.517814 |
| P43405 | Tyrosine-protein kinase SYK | SYK | | 1.650712 | 2.675003 |
| Q9NPR9 | Protein GPR108 | GPR108 | GPCRs | 3.803799 | 3.209986 |
| P28702 | Retinoic acid receptor RXR-beta | RXRB | Nuclear receptor | 1.718981 | 1.525307 |
| Q16850 | Lanosterol 14-alpha demethylase | CYP51A1 | Metabolic enzymes | 2.47602 | 2.420913 |
| O75881 | Cytochrome P450 7B1 | CYP7B1 | | 2.422192 | 1.439277 |
| Q02318 | Sterol 26-hydroxylase, mitochondrial | CYP27A1 | | 1.931162 | 1.94991 |
| P04062 | Lysosomal acid glucosylceramidase | GBA1 | | 1.552913 | 1.41248 |
| Q14534 | Squalene monooxygenase | SQLE | | 3.436055 | 2.074955 |
| P23141 | Liver carboxylesterase 1 | CES1 | | 2.066434 | 3.236823 |
| Q14849 | StAR-related lipid transfer protein 3 | STARD3 | Sterol binding proteins | 2.2104 | 2.992018 |
| Q5BJF2 | Sigma intracellular receptor 2 | TMEM97 | | 2.9527 | 1.894755 |

| | | | | | |
|---|---|---|---|---|---|
| P22307 | Sterol carrier protein 2 | SCP2 | | 2.265153 | 1.453375 |
| P35610 | Sterol O-acyltransferase 1 | SOAT1 | | 2.635481 | 1.415756 |
| P61916 | NPC intracellular cholesterol transporter 2 | NPC2 | | 1.395801 | 2.011241 |
| P08133 | Annexin A6 | ANXA6 | | 1.692172 | 1.699983 |
| P30536 | Translocator protein | TSPO | | 3.129078 | 2.078973 |
| P21796 | Voltage-dependent anion-selective channel protein 1 | VDAC1 | | 3.389149 | 4.642324 |
| P45880 | Voltage-dependent anion-selective channel protein 2 | VDAC2 | | 2.854784 | 2.11924 |
| Q9BXW6 | Oxysterol-binding protein-related protein 1 | OSBPL1A | | 2.619154 | 1.954251 |
| P22059 | Oxysterol-binding protein 1 | OSBP | | 1.239741 | 1.764535 |

**Table S2. Highlighted proteins in Fig. 1d.**

| Accession | Description | Protein | Functions | $\log_2$ Fold Change | $-\log_{10}$(P value) |
|---|---|---|---|---|---|
| Q86WV6 | Stimulator of interferon genes protein | STING | Immunology related proteins | 4.506515 | 2.764274 |
| O60603 | Toll-like receptor 2 | TLR2 | | 4.215596 | 2.398267 |
| P57764 | Gasdermin-D | GSDMD | | 5.436158 | 2.547295 |
| Q86UT6 | NLR family member X1 | NLRX1 | | 5.344272 | 3.296302 |
| P01730 | T-cell surface glycoprotein CD4 | CD4 | | 4.111781 | 2.157338 |
| Q03518 | Antigen peptide transporter 1 | TAP1 | | 4.336123 | 2.85035 |
| Q03519 | Antigen peptide transporter 2 | TAP2 | | 3.907374 | 3.216089 |
| O15533 | Tapasin | TAPBP | | 4.039073 | 2.946955 |
| Q14258 | E3 ubiquitin/ISG15 ligase TRIM25 | TRIM25 | | 5.652747 | 2.794466 |
| Q06187 | Tyrosine-protein kinase BTK | BTK | | 4.298287 | 4.112807 |
| P30273 | High affinity immunoglobulin epsilon receptor subunit gamma | FCER1G | | 6.09978 | 2.992668 |
| Q92499 | ATP-dependent RNA helicase DDX1 | DDX1 | | 3.119662 | 2.569132 |
| Q7Z2W4 | Zinc finger CCCH-type antiviral protein 1 | ZC3HAV1 | | 4.60021 | 2.664678 |
| P43405 | Tyrosine-protein kinase SYK | SYK | | 3.06876 | 2.91984 |
| P40763 | Signal transducer and activator of transcription 3 | STAT3 | | 5.114859 | 2.490776 |
| Q9NPR9 | Protein GPR108 | GPR108 | GPCRs | 5.889539 | 2.556147 |
| P41231 | P2Y purinoceptor 2 | P2RY2 | | 4.323629 | 2.133971 |
| P28702 | Retinoic acid receptor RXR-beta | RXRB | Nuclear receptor | 3.195454 | 4.031425 |
| P04150 | Glucocorticoid receptor | NR3C1 | | 4.507778 | 2.133755 |
| Q16850 | Lanosterol 14-alpha demethylase | CYP51A1 | Metabolic enzymes | 1.987532 | 2.798217 |
| O75881 | Cytochrome P450 7B1 | CYP7B1 | | 3.700157 | 2.528616 |
| Q02318 | Sterol 26-hydroxylase, mitochondrial | CYP27A1 | | 2.877427 | 1.731872 |
| P04062 | Lysosomal acid glucosylceramidase | GBA1 | | 3.979246 | 1.802669 |
| Q14534 | Squalene monooxygenase | SQLE | | 5.136105 | 2.370897 |

| | | | | | |
|---|---|---|---|---|---|
| P23141 | Liver carboxylesterase 1 | CES1 | | 4.22265 | 2.287947 |
| Q14849 | StAR-related lipid transfer protein 3 | STARD3 | Sterol binding proteins | 4.90079 | 4.202448 |
| Q5BJF2 | Sigma intracellular receptor 2 | TMEM97 | | 2.762241 | 2.036295 |
| P22307 | Sterol carrier protein 2 | SCP2 | | 3.32371 | 2.098944 |
| P35610 | Sterol O-acyltransferase 1 | SOAT1 | | 4.552104 | 1.984266 |
| P08133 | Annexin A6 | ANXA6 | | 3.930084 | 2.411957 |
| P30536 | Translocator protein | TSPO | | 6.564753 | 2.978946 |
| P21796 | Voltage-dependent anion-selective channel protein 1 | VDAC1 | | 4.439127 | 3.079632 |
| P45880 | Voltage-dependent anion-selective channel protein 2 | VDAC2 | | 5.719929 | 3.617869 |
| Q9BXW6 | Oxysterol-binding protein-related protein 1 | OSBPL1A | | 7.044447 | 2.642841 |
| P22059 | Oxysterol-binding protein 1 | OSBP | | 5.422338 | 3.074394 |

**Table S3. Highlighted proteins in Extended Data Fig. 1b.**

| Accession | Description | Protein | Functions | $\log_2$ Fold Change | $-\log_{10}$(P value) |
|---|---|---|---|---|---|
| Q86WV6 | Stimulator of interferon genes protein | STING | Immunology related proteins | 1.416057 | 2.275554 |
| O60603 | Toll-like receptor 2 | TLR2 | | 1.06993 | 1.768294 |
| P57764 | Gasdermin-D | GSDMD | | 2.953468 | 2.441102 |
| Q86UT6 | NLR family member X1 | NLRX1 | | 1.384896 | 2.707776 |
| P01730 | T-cell surface glycoprotein CD4 | CD4 | | 1.495995 | 1.761316 |
| Q03518 | Antigen peptide transporter 1 | TAP1 | | 1.581973 | 2.433191 |
| Q03519 | Antigen peptide transporter 2 | TAP2 | | 1.085646 | 2.357769 |
| O15533 | Tapasin | TAPBP | | 1.116308 | 2.448981 |
| Q14258 | E3 ubiquitin/ISG15 ligase TRIM25 | TRIM25 | | 3.271234 | 2.710326 |
| Q06187 | Tyrosine-protein kinase BTK | BTK | | 2.747415 | 2.261707 |
| Q7Z2W4 | Zinc finger CCCH-type antiviral protein 1 | ZC3HAV1 | | 1.741142 | 2.372168 |
| P40763 | Signal transducer and activator of transcription 3 | STAT3 | | 2.916002 | 2.372565 |
| Q9NPR9 | Protein GPR108 | GPR108 | GPCRs | 1.25758 | 2.031291 |
| P41231 | P2Y purinoceptor 2 | P2RY2 | | 1.667808 | 1.831363 |
| P28702 | Retinoic acid receptor RXR-beta | RXRB | Nuclear receptor | 1.26461 | 2.909963 |
| P04150 | Glucocorticoid receptor | NR3C1 | | 2.300469 | 1.995237 |
| O75881 | Cytochrome P450 7B1 | CYP7B1 | Metabolic enzymes | 1.25163 | 2.234972 |
| P04062 | Lysosomal acid glucosylceramidase | GBA1 | | 1.529378 | 1.509458 |
| Q14534 | Squalene monooxygenase | SQLE | | 1.659033 | 2.072137 |
| P04040 | Catalase | CAT | | 2.22175 | 2.62088 |
| Q14849 | StAR-related lipid transfer protein 3 | STARD3 | Sterol binding proteins | 1.72217 | 3.884437 |
| P08133 | Annexin A6 | ANXA6 | | 1.257176 | 1.991166 |
| P21796 | Voltage-dependent anion-selective channel protein 1 | VDAC1 | | 1.197229 | 2.614837 |
| P45880 | Voltage-dependent anion-selective channel protein 2 | VDAC2 | | 1.943147 | 3.322603 |

| | | | | | |
|---|---|---|---|---|---|
| Q9BXW6 | Oxysterol-binding protein-related protein 1 | OSBPL1A | | 3.027104 | 2.536851 |
| P22059 | Oxysterol-binding protein 1 | OSBP | | 3.270042 | 2.962795 |

**Table S4 Cryo-EM data collection, refinement and validation statistics**

| | HsSTING with cGAMP/C53/DCA (EMD-49094) (PDB 9N7F) |
|---|---|
| **Data collection and processing** | |
| Microscope | TFS Glacios 2 |
| Magnification | 190,000x |
| Voltage (kV) | 200 |
| Detector | TFS Falcon 4i |
| Electron exposure ($e^{-}/Å^2$) | 45 |
| Defocus range (μm) | -0.6 to -1.5 |
| Pixel size (Å) | 0.718 |
| Micrographs | 5,839 |
| Symmetry imposed | C2 |
| Final particle images (no.) | 284,997 |
| Map resolution (Å) (FSC threshold 0.143) | 2.93 |
| **Refinement** | |
| Model resolution (Å) (FSC threshold 0.5) | 3.01 |
| Map sharpening *B* factor ($Å^2$) | -100.9 |
| Model composition | |
| Non-hydrogen atoms | 10,172 |
| Protein residues | 1270 |
| Ligands | 4 |
| Map CC | 0.83 |
| R.m.s. deviations | |
| Bond lengths (Å) | 0.003 |
| Bond angles (°) | 0.558 |
| Validation | |
| MolProbity score | 1.45 |
| Clashscore | 8.22 |
| Poor rotamers (%) | 0.00 |
| Ramachandran plot | |
| Favored (%) | 99.11 |
| Allowed (%) | 0.89 |
| Disallowed (%) | 0.00 |

### Bile acid photoaffinity probe characterization

The $^{1}$H-NMR and $^{13}$C-NMR spectra were recorded on a Bruker 600 MHz spectrometer running TopSpin 2.1 in indicated deuterated solvents using TMS or solvent peak as a standard. All $^{13}$C-NMR spectra were recorded with complete proton decoupling. TopSpin (4.0.7) was used for NMR spectrum analysis. The synthetic routes were adapted from ref. 35.

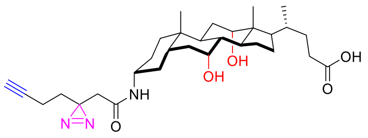

**(R)-4-((3R,5S,7R,8R,9S,10S,12S,13R,14S,17R)-3-(2-(3-(but-3-yn-1-yl)-3H-diazirin-3-yl)acetamido)-7,12-dihydroxy-10,13-dimethylhexadecahydro-1H-cyclopenta[a]phenanthren-17-yl)pentanoic acid (CA2).** $^{1}$H-NMR (600 MHz, $CDCl_3$, representative signals) δ (ppm) 6.17 (d, $J$ = 8.2 Hz, 1H), 4.00 (s, 1H), 3.86 (s, 1H), 3.67-3.57 (m, 1H), 2.44-2.35 (m, 1H), 2.08 (s, 1H), 0.99 (d, $J$ = 5.9 Hz, 3H), 0.90 (s, 3H), 0.69 (s, 3H). $^{13}$C-NMR (151 MHz, $CDCl_3$) δ 178.59, 167.20, 82.98, 73.30, 69.88, 68.67, 49.92, 46.95, 46.57, 42.08, 41.81, 41.63, 39.53, 36.42, 35.79, 35.34, 34.76, 34.55, 32.15, 30.95, 30.89, 28.30, 27.80, 27.65, 26.76, 26.24, 23.28, 22.76, 17.44, 13.43, 12.62. HRMS (ESI) calculated for $C_{31}H_{47}N_3O_5$ $[M+H]^+$ : 542.3594, found 542.3573.

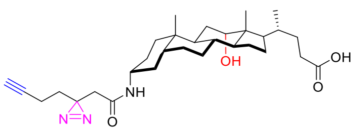

**(R)-4-((3R,5R,8R,9S,10S,12S,13R,14S,17R)-3-(2-(3-(but-3-yn-1-yl)-3H-diazirin-3-yl)acetamido)-12-hydroxy-10,13-dimethylhexadecahydro-1H-cyclopenta[a]phenanthren-17-yl)pentanoic acid (DCA2).** $^{1}$H-NMR (600 MHz, $CDCl_3$, representative signals) δ 5.95 (d, $J$ =

8.3 Hz, 1H), 4.01 (s, 1H), 3.84-3.72 (m, 1H), 2.44-2.35 (m, 1H), 2.12 (t, $J$ = 2.7 Hz, 1H), 0.97 (d, $J$ = 6.4 Hz, 3H), 0.92 (s, 3H), 0.68 (s, 3H). $^{13}$C-NMR (151 MHz, $CDCl_3$) δ 179.15, 167.04, 82.90, 73.56, 70.00, 49.75, 48.51, 47.44, 46.64, 42.35, 41.79, 36.05, 35.80, 35.18, 34.18, 33.77, 33.45, 32.05, 31.08, 30.83, 28.60, 27.69, 27.60, 26.96, 26.23, 26.04, 23.74, 23.31, 17.49, 13.44, 12.86. HRMS (ESI) calculated for $C_{31}H_{47}N_3O_4$ $[M+H]^+$ : 526.3645, found 526.3640.

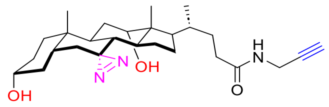

**(R)-4-((3R,5R,8R,9S,10S,12S,13R,14S,17R)-3,12-dihydroxy-10,13-dimethyl-1,2,3,4,5,6,8,9,10,11,12,13,14,15,16,17-hexadecahydrospiro[cyclopenta[a]phenanthrene-7,3'-diazirin]-17-yl)-N-(prop-2-yn-1-yl)pentanamide (DCA4).** $^{1}$H-NMR (600 MHz, $CDCl_3$, representative signals) δ (ppm) 5.72 (s, 1H), 4.05-4.01 (m, 2H), 3.96 (s, 1H), 3.64-3.56 (m, 1H), 2.22 (t, $J$ = 2.6 Hz, 1H), 1.00 (s, 3H), 0.95 (d, $J$ = 6.3 Hz, 3H), 0.61 (s, 3H), 0.02 (d, $J$ = 14.5 Hz, 1H). $^{13}$C-NMR (151 MHz, $CDCl_3$) δ 173.09, 79.82, 72.23, 71.69, 71.45, 47.29, 45.95, 43.30, 41.89, 37.85, 37.56, 35.22, 35.01, 34.83, 34.79, 33.19, 32.93, 31.47, 30.39, 30.30, 29.31, 28.49, 27.32, 24.42, 22.51, 17.72, 12.81. HRMS (ESI) calculated for $C_{27}H_{41}N_3O_3$ $[M+H]^+$ : 456.3226, found 456.3214.

**Data S1.** Proteomics dataset for bile acid reporter-enriched proteins in THP-1 cell line.

**Data S2.** Cellprofiler pipeline used for STING-GFP puncta analysis.
